# Inositol Pyrophosphates Mediate Chloroplast Lipid Remodeling and Nuclear Gene Repression during High-Light Acclimation in *Chlamydomonas reinhardtii*

**DOI:** 10.64898/2026.06.04.730061

**Authors:** Rodrigo Bedera-García, Luis G. Heredia-Martínez, M.E. García-Gómez, Borja Prieto-Muñiz, José M. Ortega, Inmaculada Couso

**Affiliations:** Microalgae Systems Biology and Biotechnology Research Group, Institute for Plant Biochemistry and Photosynthesis, Spanish National Research Council; Departamento de Bioquímica Vegetal y Biología Molecular, Facultad de Biología, Universidad de Sevilla, 41012 Seville, Spain; Instituto de Bioquímica Vegetal y Fotosíntesis, cicCartuja, Universidad de Sevilla and CSIC, 41092 Seville, Spain

**Keywords:** Inositol polyphosphates, Chlamydomonas, RNAseq, photosynthesis regulation and cell signaling

## Abstract

Microalgae are photosynthetic organisms capable of autotrophic growth. Their applicability in multiple industrial fields has been largely studied, thanks to their ability to fixate CO_2_ into high added value organic products like fatty acids and carotenoids. However, our understanding of the cellular signaling networks that control carbon flux and acclimation to environmental stress remains incomplete. In this study, we used the *Chlamydomonas reinhardtii* mutant strain *vip1-1*, which carries a loss-of-function mutation in the hexakisphosphate kinase responsible for the synthesis of inositol pyrophosphates InsP_7_ and InsP_8_ (PP-InsPs), to investigate the role of these molecules during high-light acclimation. Our results indicate that PP-InsPs participate in the regulation of carbon storage in the form of starch and their deficiency increases TAGs levels in the algal cells. They also impact chloroplast-specific lipid remodeling by modifying membrane composition and fluidity through fatty acid desaturations and glycerolipid composition. In addition, our findings suggest that PP-InsPs are involved in chloroplast-nucleus communication, where they coordinate transcriptional repression of photosynthesis associated nuclear genes (PhANGs), fatty acid desaturases and lipid synthases, contributing to cellular acclimation to high light. We also found that PP-InsPs modulating effect extended to protein synthesis and accumulation of Calvin-Benson-Bassham cycle intermediates. Therefore, we propose that PP-InsPs function as integratory molecules that balance carbon allocation between storage and structural pools, in response to environmental cues such as high light. These data uncover a novel function of PP-InsPs in high light acclimation and potentially in chloroplast-nucleus communication, providing new insights that may help engineering more resilient and efficient strains.

## 1 Introduction

Understanding responses and adaptability of green organisms to changeable conditions are key steps towards biotechnological applications and strain improvements in microalgae. In this way, central controllers and signaling molecules are important integrators of external signals and need to be evaluated as potential targets for applied studies. Inositol polyphosphates are a well conserved family of signaling molecules in eukaryotes, produced through sequential phosphorylation of the six-carbon inositol ring. The synthesis of inositol polyphosphates starts from the de-phosphorylated molecule *myo*-inositol. This compound is used as a substrate of consecutive kinases to produce phytic acid, also known as InsP_6_ (Williams et al., 2015; Desfougères et al., 2019; Cridland and Gillaspy, 2020). In the model green alga *C. reinhardtii*, one of the two classes of inositol pyrophosphate synthetases found in eukaryotes is conserved, which is the diphosphoinositol pentakisphosphate kinase VIP1. InsP_6_ is predicted to undergo two alternative pyrophosphorylations which result in 1-diphosphoinositol 2,3,4,5,6-pentakisphosphate (1-PP-InsP_5_) catalyzed by VIP1 or 5-diphosphoinositol 1,2,3,4,6-pentakisphosphate (5-PP-InsP_5_) (both InsP_7_), and combined in 1,5-bis-diphosphoinositol 2,3,4,6-tetrakisphosphate (1,5PP-InsP_4_) (InsP_8_) (Williams et al., 2015; Laha et al., 2015, Laha et al., 2021). However, more information is needed regarding the positions of the pyrophosphates in *Chlamydomonas* as more recently the isomer 4/6-InsP_7_ has been identified in other green organisms such as *A. thaliana* and *M. polymorpha* (Laha et al., 2022; Chalak et al., 2024; Laurent et al, 2024; Pullagurla et al., 2025, Yadav et al., 2025), which indicate the complexity of PP-InsP signaling network employed by green organisms. In this sense, these pyrophosphorylated species are usually referred to as inositol pyrophosphates, or PP-InsPs.

These PP-InsPs, among all inositol polyphosphate species, have received the most attention in plants, yeasts and mammals due to their regulatory functions. Traditionally, the main role attributed to these molecules has been the intracellular P homeostasis, which has been detected in plants like *A. thaliana*, *Oryza sativa* or *Marchantia polymorpha* (Wild et al. 2016, Dong et al. 2019, Wang et al. 2021, Guan et al. 2022, Pullagurla et al. 2025), yeasts (Gu et al. 2017, Chabert et al. 2023, Pipercevic et al. 2023) or even humans (Azevedo and Saiardi 2017, Zuo et al. 2025). Nevertheless, new functions of PP-InsPs have been reported in the green lineage. In *Arabidopsis thaliana*, they have been found participating in hormone response complexes, such as the auxin receptor complex (Laha et al. 2022) or the jasmonate coreceptor complex (Laha et al. 2015). Additionally, a role for PP-InsPs in nitrate homeostasis and cell wall biogenesis has been proposed recently in *Marchantia polymorpha* (Laurent et al. 2024), and the proteins putatively responsible for 4/6-InsP_7_ production (IPMK homologs) have been found to be involved in heat stress responses in *Arabidopsis thaliana* and *Marchantia polymorpha* (Yadav et al. 2025). In microalgae, the inositol polyphosphate profile has been reported to respond to CO_2_ levels in *Chlorella*, suggesting a role in carbon assimilation in unicellular photosynthetic organisms (Morales-Pineda et al. 2023). Furthermore, we recently uncovered in *C.reinhardtii* a new function of the PP-InsPs in NIT2-dependent nitrogen starvation responses (Bedera-García et al. 2025). These reports indicate that the signaling roles of PP-InsPs and the cellular processes they influence remain incompletely understood, particularly in the green lineage, where new functions of these small regulators continue to be uncovered.

In 2016, a *C. reinhardtii* mutant named *vip1-1* was identified during a hypersensitivity screening using rapamycin (TOR inhibitor) (Couso et al., 2016). *vip1-1* was found to carry a loss-of-function mutation in the gene encoding VIP1 (an inositol polyphosphate kinase) and it was demonstrated that PP-InsPs affect several processes controlled by TOR regulatory pathway, including carbon metabolism, autophagy and nutrient sensing (Couso et al. 2016, Couso et al, 2021). This mutation resulted in a very strong reduction of PP-InsPs levels that affected carbon flux related functions like tricarboxylic acid cycle and TAG accumulation (Couso et al. 2016, Couso et al. 2021). The phosphoproteome analysis of *vip1-1* uncovered multiple altered targets in photosynthetic proteins, with a remarked deregulation in the phosphosites of light-harvesting complex II (LHCII) proteins (Couso et al., 2021) indicating differences in carbon assimilation and photosynthesis performance that may affect the adaptability to light stress.

The high light response in photosynthetic organisms enables survival when irradiance exceeds the light-harvesting capacity of the photosynthetic machinery. *C. reinhardtii* has been used extensively as a model organism to investigate stress response mechanisms including high light response, owing to the simplicity of being unicellular while sharing many regulatory pathways and conserved proteins with plants (Bonente et al. 2012, Allorent et al. 2013, Erickson et al. 2015, Dupuis et al. 2025). In this regard, the investigation of PP-InsPs signaling in combination with the response to high irradiances is of great interest not only for microalgal biotechnology, but also for its potential relevance to other photosynthetic organisms. Therefore, to improve our knowledge on the possible roles of PP-InsPs in high light signaling, we conducted an integrative study involving a metabolomic and transcriptomic analysis of wild-type and *vip1-1* strains under standard irradiances and high light conditions. Our results indicate that PP-InsPs are involved in the high light acclimation response through the coordination of carbon allocation between storage and structural pools, and through adjusting thylakoid membrane fluidity. Additionally, here we report that these signaling molecules take part in a photosynthesis regulation through transcriptional repression. Taken together, these results position inositol pyrophosphates as signaling molecules that integrate light cues with carbon sink management and gene expression to adjust functional and structural remodeling of the chloroplast.

## 2 Materials and Methods

### 2.1 Strains, culture conditions and growth curves

*Chlamydomonas reinhardtii* strain CC-1690 wild-type MT+ (Sager 21gr) (Sager, 1955) was used as control and compared to *vip1-1*, an insertional mutant of the CC-1690 strain (Couso et al., 2016). The experiments were performed at exponential growth phase (1-2 × 10^6^ cells ml^−1^) in liquid TAP (tris acetate phosphate) medium at 25 °C, where the control light conditions (standard light, SL) were set at 50 μE m^−2^ s^−1^ and the high light conditions (HL) were set at 1500 μE m^−2^ s^−1^, over a period of 24 h. Growth curves were performed using OD750nm measurements under the two light conditions, starting with SL until 24h and then subjected to HL in the WT, *vip1-1* and complemented *vip1-1:VIP1*.

### 2.2 Chlorophyll determination and thermoluminescence

Chlorophyll determination was done as described in Bedera-García et al. (2025). Two independent biological replicates and two technical replicates were used for chlorophyll determination and activity assays. 1×10^6^ cells mL^−1^ cultures of WT and *vip1-1* grown under standard or high light were assayed. Chlorophyll determination was performed collecting cells by centrifugation at 6500 *g*, resuspended in methanol and incubated at 70 °C for 15 min. The supernatant obtained was then centrifugated at 6500 *g* and used to perform OD determination using an Amersham Biosciences Ultrospec 3100 pro spectrophotometer. The concentration of chlorophyll in the different samples was then calculated by measuring the absorbance at 650 and 665 nm, according to Mackinney (1941).

Thermoluminescence (TL) glow curves of *C. reinhardtii* cell suspensions were obtained using a home-built apparatus designed by Dr. Jean-Marc Ducruet for luminescence detection from 1 °C to 80 °C. A detailed description of the system can be obtained elsewhere (Ducruet 2003; Roncel et al. 2016; García-Calderón et al. 2019; Castell et al. 2021). Briefly, temperature regulation, signal recording and flash sequences were driven by a computer through a National Instrument DAQ-Pad 1200 interface, using a specific acquisition program developed by Dr. Ducruet. The sample cuvette consisted in a horizontal chamber (2 cm diameter) with a copper film on the bottom. An electrically insulated resistor heater (Thermocoax), powered by a variable (0 to 5 A) computer-driven power supply, was mounted below the chamber for temperature regulation. The resistor heater was cooled by a temperature-controlled bath. Luminescence emission was detected by a H5701-50 Hamamatsu photomultiplier module. Illumination was performed through a light guide parallel to the photomultiplier, both of them being attached to the same stand sliding horizontally from the illumination to the measuring position. Single turn-over flashes were provided by a xenon white light (WalzXST-103). TL measurements were carried out as reported in Roncel et al. (2016). Typically, 200 ml *C. reinhardtii* cell suspensions were dark-incubated for 2 min at 20 °C, then cooled to 1 °C for 2 min. After this period, cells were illuminated with two saturating single-turnover flashes separated by 1 s. Luminescence emission was then recorded while warming the samples from 1 °C to 65 °C at a heating rate of 0.5 °C per second. TL parameters were obtained from three independent measurements. Data acquisition, signal analysis and graphical simulation were performed as previously described (Ducruet and Miranda 1992; Zurita et al. 2005; Ducruet et al. 2011). Most of the experiments were performed using suspensions with a cell concentration of ≈2.4 × 10^7^ cells mL^−1^.

### 2.3 Inositol polyphosphates analysis

Inositol polyphosphates were extracted using 5% TCA and analyzed by LC-MS/MS as reported in Couso et al., 2016. PP-InsPs (InsP_7_ and InsP_8_) data were analyzed using the QualBrowser and QuanBrowser applications of Xcalibur (Thermo Fisher Scientific). Data were normalized using the internal standard 3-fluoro-InsP3. Mean data and SD were calculated from three biological replicates and three technical replicates.

### 2.4 Transmission electron microscopy

Cultures were collected at 1-2×10^6^ cells mL^−1^ and fixed with 2.5% glutaraldehyde in 0.1 M Na-cacodylate buffer at pH 7.4. Samples fixation and sections were obtained following the same procedure as reported in Bedera-García et al., 2025. Sections were examined in a Zeiss Libra 120 transmission electron microscope and digitized (2048 × 2048 × 16 bits) using an on-axis mounted TRS camera. WinTEM™ was used to analyze images.

### 2.5 Lipid fractionation and analysis

Total lipids were extracted from *C. reinhardtii* cells grown under standard and high light conditions, following the method described by Díaz-Santos et al. (2025) with some modifications. Briefly, 0.1-0.3 g of dry cell weight (DCW) of *C. reinhardtii* cells were resuspended in chloroform:methanol:water (1:2:2) and disrupted by ultrasonic treatment for 20 min. The lower chloroform layer containing the lipids was collected. Re-extraction process of the upper layer was performed to improve the lipid extraction method. Finally, the solvent was completely removed using N_2_ and total lipids were dissolved in chloroform for further analysis. Polar lipids analysis was performed by total lipids separation using thin layer chromatography (Hernández et al. 2008). Individual lipids were visualized by iodine vapor, and their identification was carried out by comparing them with reference standard, Fatty acid methyl esters of total lipid extraction and the individual polar lipid classes were produced by acid-catalyzed transmethylation (Garcés et al., 2023) and analyzed by gas chromatography using a GC-MS-QP2010 Plus (Shimazu, Kyoto, Japan) (Gallardo-Martínez et al., 2023). Heptadecanoic acid was used as an internal standard to calculate fatty acid contents in the samples. The results obtained are presented as means (µg mg of DCW^−1^) ± SD. Significance was calculated using t-test.

### 2.6 Starch Quantification

Starch granules in dry pellets were alkaline solubilized with 1mL of 0.2M KOH and heated at 100°C. After 30 min, samples were gradually cooled and pH was adjusted to 5.0 adding 300μL of 1M acetic acid. To start digestion, 7.4U of α-amylase were added and incubated 30 min at 37°C, breaking down starch in small linear and branched oligosaccharides. After that, 5U od amyloglucosidase were added and incubated 1-2 h at 55°C releasing glucose residues. Finally, in order to stop enzymatic reaction, the samples were incubated at 100°C during 2 min and centrifuged at 13000 xg for 10 min discarding pellet. Enzymes were prepared in 0.1 M of sodium acetate pH 4.5.

### 2.7 Transcriptomic data generation and analysis

For RNA-seq data, three independent biological replicates were considered for each strain and condition: WT and *vip1-*1 strains grown under standard and high light. RNA extraction was performed using mechanical disruption of the frozen cell pellets in a Mini Bead Beater (Biospe Products) mixed with 0.5 mm glass beads for individual cell lysis in the presence of an extraction buffer consisting of phenol:chloroform (1:1 v/v). RNA was purified using the ISOLATE II RNA Plant Kit (Bioline) following the manufacturer’s instructions. The quality of the RNA was first analyzed by gel electrophoresis, and the RNA integrity number (RIN) was computed using Agilent 2100 Bioanalyzer. RNA-seq data generation was obtained as described in Serrano-Pérez et al. (2022) and analyzed with the help of the MARACAS tool (Romero-Losada et al., 2022). GO and KEGG enrichments were computed using genes differentially expressed, with a threshold of fold change > 2 and adjusted p-value < 0.05. Approximately 30 million 100 nt reads were obtained per sample. The RNA-Seq data was processed using fastQC (https://www.bioinformatics.babraham.ac.uk/projects/fastqc/) to perform the quality control, Hisat2 (Kim et al., 2019) to map reads, Stringtie (Pertea et al., 2015) to assemble the transcripts and Ballgown (Frazee et al., 2015) to quantify expression. Raw data was normalized using the upper quantile method. Differential expression analysis was performed using the Limma R package (Ritchie et al., 2015) and the BH method was used to adjust *p*-values. Target location of the gene product was obtained using PredAlgo annotation (Tardif et al. 2012), and the “Other” category was assumed to be cytosolic, based on the known location of gene products contained in this category.

### 2.8 RT-qPCR data generation

The gene expression of chloroplast encoded genes was assayed through RT-qPCR. cDNA was generated using QuantiTect Reverse Transcription Kit (205311, Qiagen) and assayed using SensiFast SYBR Fluorescein Kit (BIO-96020, Meridian Bioscience) and the CFX96 real-time detection system (Bio-Rad). After heating at 95°C for 2 minutes, 40 cycles of 95°C for 5 s, 55°C for 10 s and 72°C for 20 s were done. The specificity of the amplification was confirmed by performing a melting curve from 65°C to 95°C with 0.5°C/min increments. Relative abundance of cDNA was calculated with the ΔΔCt method using *cblp* as a housekeeping gene (von Kampen et al. 1994). The primers used to analyse each gene expression were: *cblp* left primer CACCCAGTCCTCCATCAAGA, *cblp* right primer TGGGCATTTACAGGGAGTGG; *psbA* left primer ATGGGTCGTGAGTGGGAATT, *psbA* right primer GAACCACCGAATACACCAGC; *psbB* left primer AGCATTAGGTCAACGTCGTT, *psbB* right primer TCACCAATTGTACCACCAGC; *psbD* left primer ATTCTGGGCAGCTTTCGTTG, *psbD* right primer GATGCGTGACCTAACCAACC and *chlN* left primer AATCTTGTCCACAACGTCGT, *chlN* right primer ACTCGACCACCACGTACTAA. At least, three technical replicates were done for each independent biological replicate.

### 2.9 Metabolomic data generation and analysis

Four independent biological replicates were assessed for each strain and condition. The metabolite content determination was performed using cell pellets that were first lyophilized (Skadi-Europe TFD 8503) and stored at −20°C as described in Serrano-Pérez et al. (2022). Metabolites were determined from 15-20 mg of lyophilized cell biomass subjected to mechanical disruption in a Mini Bead Beater (Biospe Products) with 0.5 mm glass beads in the presence of 1 mL extraction buffer consisting of chloroform:methanol (3:7 v/v). Centrifugation at 5000 g for 5 min at RT was then performed and these two steps were repeated until the pellets were colorless. These pellets are then resuspended in MiliQ water to separate aqueous phases that were concentrated using a speed-vacuum (Eppendorf concentrator Plus). Samples were then resuspended using 200 µL of MiliQ water and submitted to UPLC/MS analysis as described in McCloskey and Ubhi (2015). Raw data was normalized using 10 µM paracetamol as an internal standard and the dry weight of each sample. The significance was calculated using the Wilcoxon signed-rank test. For heatmap plotting, metabolites were auto-scaled and the rows were clustered using euclidean distance and “complete” method from stats R package (R Core Team, 2023).

### 2.10 Carotenoid determination and quantification

Three independent biological replicates for each condition were analyzed. Cells were collected by centrifugation at 6500 *g*. Pellets were resuspended in 0.5 mL TAP media and and heated at 90°C for 3 minutes. Then, 2 mL of acetone were added. The mixture was centrifuged at 10000 *g* for 15 minutes. The supernatant containing the carotenoids was collected, and the solvent was evaporated using a N_2_ atmosphere. Samples were resuspended in 600 µL acetone for analysis. The determination and quantitation were done in HPLC (Hitachi Elite LaChrom) with a photodiode-array detector (Hitachi L-2455). Separation was achieved using a Waters NovaPak C-18 (3.9 x150 mm, 4 µm particle size, 60 Å pore size) with the following eluents: eluent A (0.1 M ammonium acetate and 15:85 v/v H_2_O:methanol) and eluent B (44:43:13 v/v methanol:acetonitrile:acetone) using the program reported in Böhme et al. 2002. Temperature was kept at 20°C and flow was set to 800 µL min^−1^.

## 3 Results

### 3.1 PP-InsPs deficient mutant shows important sensitivity to HL stress

To interrogate whether PP-InsPs influence acclimation to high light exposure, we examined a growth curve of the green model alga *Chlamydomonas reinhardtii vip1-1* strain which is PP-InsPs-deficient (Couso et al. 2016) and its corresponding parental (WT) and complemented strains under standard light (SL) and high light (HL) conditions (Figure 1A). We subjected cultures to HL after 24h cultivated under SL to monitor specific differences beyond this time point (Figure 1A). We observed important differences in growth in *vip1-1* after 8 hours of HL exposure (36 hours after the starting point) (Figure 1A) reaching 30% less cell density after 60 hours in *vip1-1* than in the WT, indicating sensitivity of the culture to these stress conditions.

**Figure 1:**
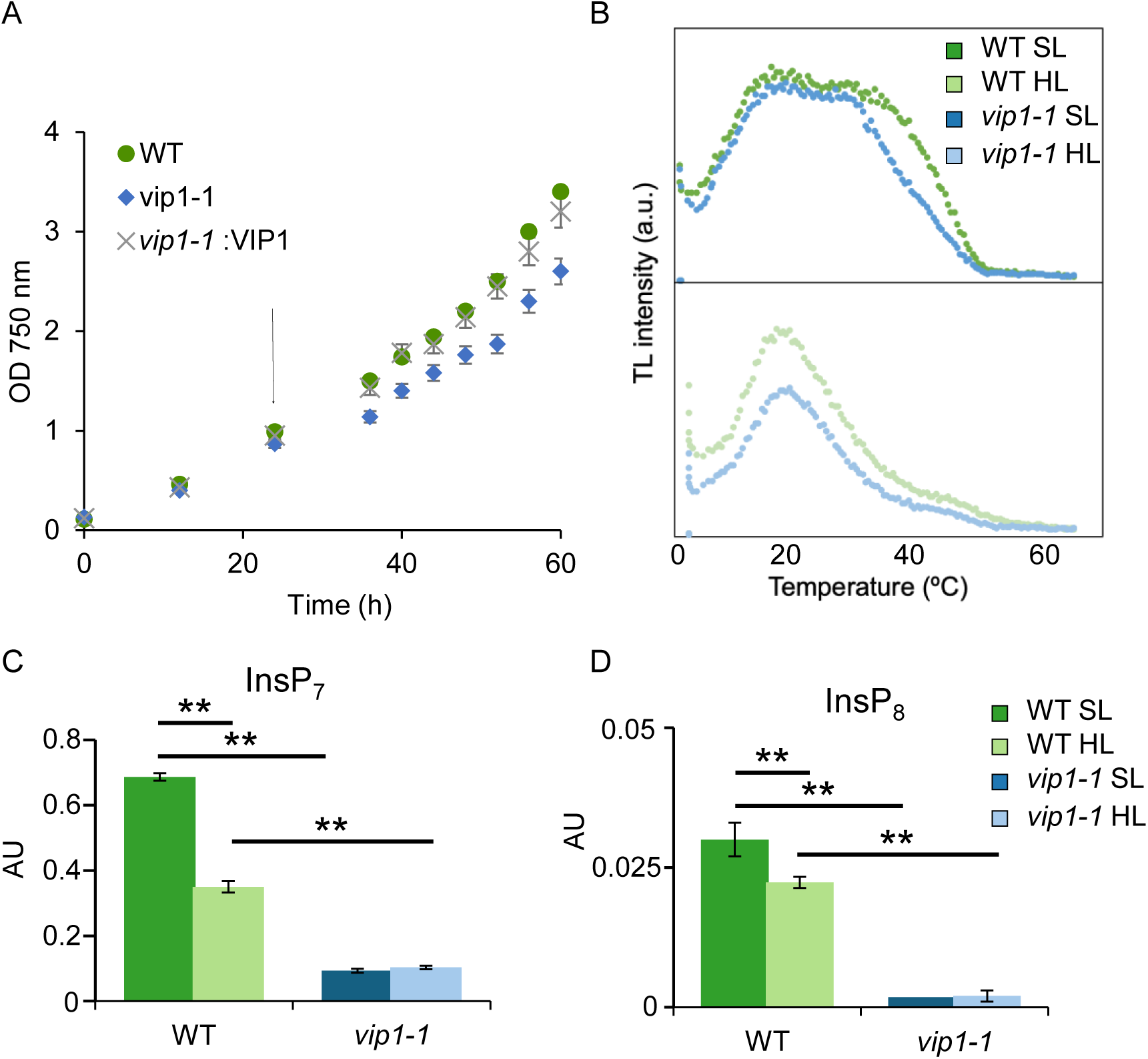
(A) Growth curve of WT, vip1-1 and the complemented strain vip1-1 :VIP1 under high light conditions. Arrow indicates the starting point of HL stress. (B) Representative standard TL glow curves of WT and vip1-1 strains cultured under control (upper panel) and high light illumination conditions (bottom panel). (C) InsP7 and (D) InsP8 levels of WT and vip1-1 under SL and HL.

The effect of *vip1-1* insertion on the electron transfer activity of PSII in *C. reinhardtii* cells was also studied using the thermoluminescence (TL) technique. First, we studied some of the most relevant TL emission bands in *C. reinhardtii* cell from WT and *vip1-1* cell lines cultured under control conditions (SL) (Figure 1 B, upper panel). Excitation of WT and *vip1-1* dark-adapted cell suspensions with two saturating single turn-over flashes at 1 °C induced the appearance of very complex TL glow curves. These TL signals could be well simulated by three decomposition components with different temperatures of the maximum intensity (t_max_) and contributions to the total signal intensity. We assigned the two first components to the well-known TL B1 and B2 emission bands originating from the recombination reactions of S_3_Q_B_^−^ and S_2_Q_B_^−^ charge pairs, respectively. The third component appearing at higher temperatures was assigned to the so-called afterglow (AG) TL emission band (Miranda and Ducruet 1995). This TL emission seems to reflect a back-flow of electrons from unknown reductants present in the stroma to the quinone acceptors of PSII, allowing their recombination with S_2_ and S_3_ states (Sundblad et al. 1988; Miranda and Ducruet 1995; Havaux 1996). The mathematical analysis of these three components allowed us to estimate t_max_ and contributions of B and AG bands to the total signal intensity. For the B1 band, we obtained t_max_ values of about 15°C for both WT and *vip1-1*, and signal contributions of 24% (WT) and 20% (*vip1-1*). For the B2 band, we obtained t_max_ values of about 25 °C for both WT and *vip1-1*, and signal contributions of 32% (WT) and 47% (*vip1-1*). For the AG band, t_max_ values of about 39 °C for both WT and *vip1-1* and signal contributions of 44% (WT) and 33% (*vip1-1*) were obtained. Thus, under control conditions, the main effect observed of the *vip1-1* insertion was a significant decrease (about 25%) of the contribution of the AG band to the total TL intensity, and consequently of the ratio between the signal intensities of the AG and B bands (from 0.79 in WT to 0.49 in *vip1-1*). No significant changes were detected in t_max_ values of B and AG bands for WT and *vip1-1* lines. The intensity of the AG band is an indicator of the assimilatory potential (NADPH+ATP) when it is inducible by a white light illumination (Mellvig and Tillberg 1986; Palmqvist et al. 1986; Sundblad et al. 1986) or xenon flash excitation (Krieger et al. 1998). The very significant decrease in the ratio between the signal intensities of the AG and B bands observed in *vip1-1* cells in comparison with WT cells under control conditions may be due to a lower availability of NADPH and/or ATP.

We have also analyzed TL emission bands of *C. reinhardtii* WT and *vip1-1* cell lines grown under HL (Figure 1B, bottom panel). The TL glow curves obtained after excitation of dark- adapted cell suspensions with two saturating single turn-over flashes at 1 °C could be also well simulated by three decomposition components: B1, B2 and AG bands. WT and *vip1-1* did not show significant differences in the parameters of the B and AG bands. The t_max_ and intensities contribution values obtained for the two cell lines were, respectively, 16 °C and 49% (B1 band), 25 °C and 33% (B2 band), and 39 °C and 18% (AG band). However, a significant decrease in the total TL signal intensity (of ≈25%) was observed for *vip1-1* compared to WT cells. The amplitude of the TL signal is related to the overall PSII activity from the water-splitting system to the final quinone acceptor (Rutherford et al 1984; Msilini et al. 2011). The significant TL signal drop observed in *vip1-1* cells suggests a decrease in the number of photochemically active PSII complexes under HL conditions (Msilini et al 2011).

We also analyzed PP-InsPs levels in both strains using LC-MS/MS following the protocol reported in Couso et al., 2016 under SL and HL conditions (Figure 1C and D). The levels of InsP_7_(Figure1C) and InsP_8_ (Figure1D) in WT were significantly downregulated under HL conditions showing 48% decrease in InsP_7_ and 33% decrease in InsP8 while no significant differences were observed in the mutant as expected. These results suggest that the synthesis of PP-InsPs is specifically responsive to HL, warranting further investigation into how these molecules contribute to cellular regulation under stress.

### 3.2 TEM analysis shows a deregulation of carbon storage in *vip1-1* cells under HL stress

Ultrastructure analysis has previously indicated important differences in *vip1-1* cells under nutritional stress (Bedera-Garcia et al., 2025). In this regard, we performed TEM analysis of WT and *vip1-1* under SL and HL stress to visualized potential differences among these samples. We found that *vip1-1* cells showed reduced levels of starch granules especially under HL conditions when compared with WT. In fact, the number of starch granules in WT cells after HL stress highly increased while *vip1-1* cells did not (Figure 2A). In order to confirm these observations, we measured starch levels in the different samples and found a significant accumulation of starch under HL in WT (Figure 2B). In contrast, *vip1-1* cells did not upregulate starch levels and the quantitation showed no significant differences between SL and HL conditions in the mutantś samples (Figure 2B). These results indicate an aberrant regulation of carbon storage that we further analyzed by quantifying total FAs (Figure 2C) and TAGs (Figure 2D). We found around 15% and 25% higher levels of total FAs in *vip1-1* under SL and HL respectively compared to WT while we found a significant decrease in WT under HL conditions (Figure 2C). Additionally, we used lipid fractionation approach and performed FAMEs identification and quantification using GC-MS. We found significant differences in all the fatty acids species analyzed between WT and *vip1-1* but C16:0, C16:3, C16:4 and C18:3 levels were especially different under HL (Supplemental Figure 1). After lipid fractionation, we also analyzed TAGs fraction and found an important upregulation of these lipid class in *vip1-1* samples reaching 2.25 levels of the WT samples under HL (Figure 2D). These results confirmed an important deregulation of lipid metabolism that impacts C storage in PP-InsPs deficient cells that needed further exploration.

**Figure 2:**
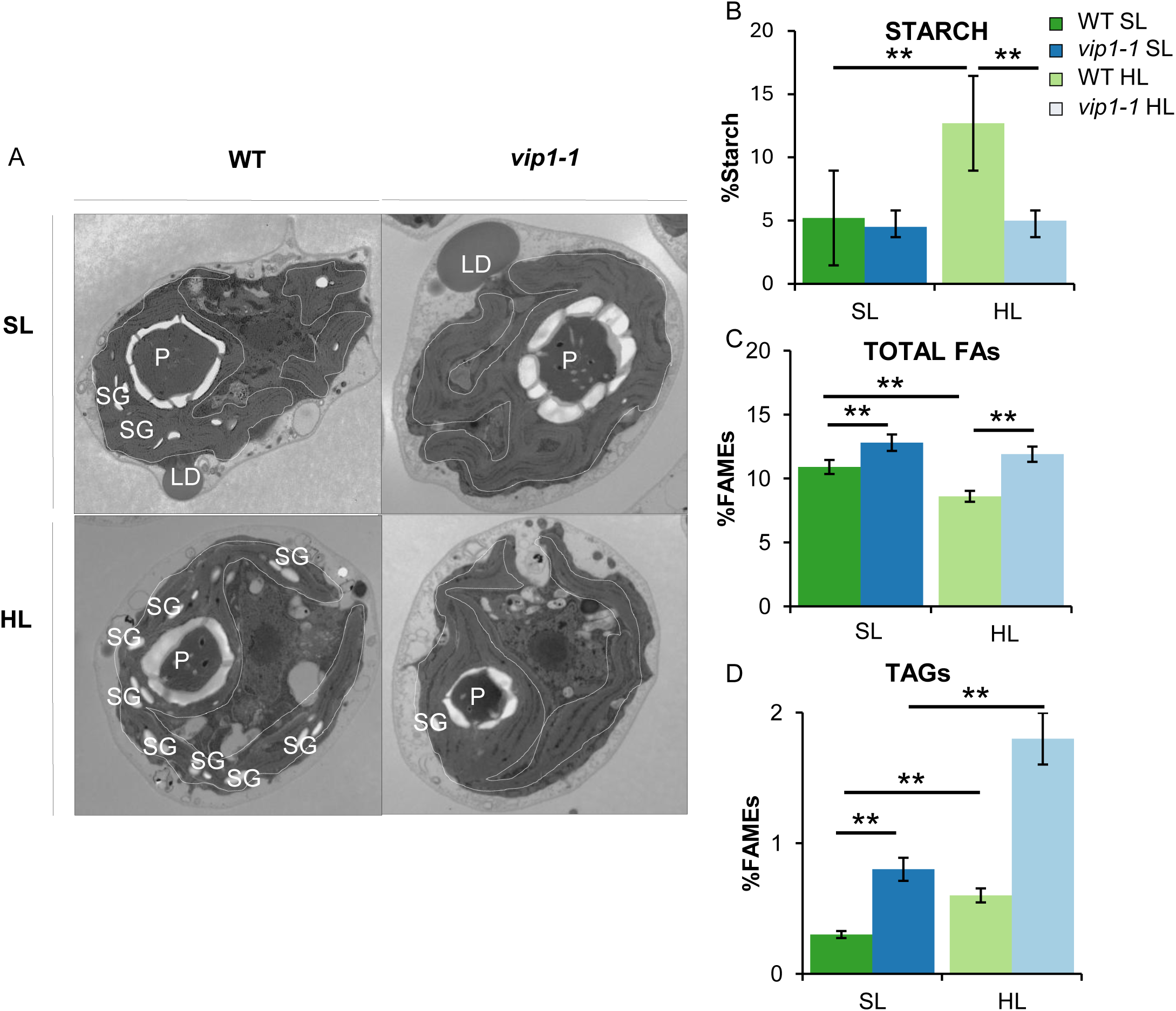
(A) Ultrastucture analysis (TEM) of WT and vip1-1 under SL and HL; SG: starch granules; P pyrenoid ; LD: lipid droplet. (B) Starch levels in WT and vip1-1 under SL and HL (C) Total lipids (FAMES) in WT and vip1-1 under SL and HL (D) TAGs levels in WT and vip1-1 under SL and HL.

### 3.3 Lipid fractionation in *vip1-1* samples indicate remodeling of lipid synthesis under HL

Previous studies have shown that PP-InsPs regulate lipid metabolism and photoprotection (Couso et al. 2016, Couso et al. 2021). Additionally, structural lipids present in the thylakoid membrane such as monogalactosyldiacylglycerol (MGDG), digalactosyldiacylglycerol (DGDG), sulfoquinovosyldiacylglycerol (SQDG) and phosphatidylglycerol (PG) have been tightly connected to photosynthetic function, and defective mutants on these lipids have shown compromised photosynthesis in different green organisms (Gombos et al. 2002; Laczkó-Dobos et al. 2008, Kobayashi et al. 2013; Fujii et al. 2014; Zhu et al. 2024). Therefore, we conducted lipid analysis to evaluate whether deregulation of structural tylakoid lipids might be the source of impaired photosynthetic efficiency. In this study, we analyzed different lipid classes, galactolipids (MGDG and DGDG), SQDG and phospholipids (PG/PE). Our results indicate that *vip1-1* cells growing under SL conditions significantly accumulated less MGDG, SQDG and PG/PE (2.96, 1.80 and 2.27-fold respectively) (Figure 3A). However, under HL stress, the two strains exhibited opposed responses. The WT strain significantly lowered MGDG and PG/PE levels in HL (1.66 and 1.73-fold respectively) compared to SL conditions, whereas the *vip1-1* strain significantly accumulated higher amounts of MGDG, DGDG, SQDG and PG/PE in the same conditions (2.16, 2.24, 1.50 and 4.33- fold) (Figure 3A). Overall, *vip1-1* strain overaccumulates all the detected polar lipid species under HL compared to the WT strain.

**Figure 3:**
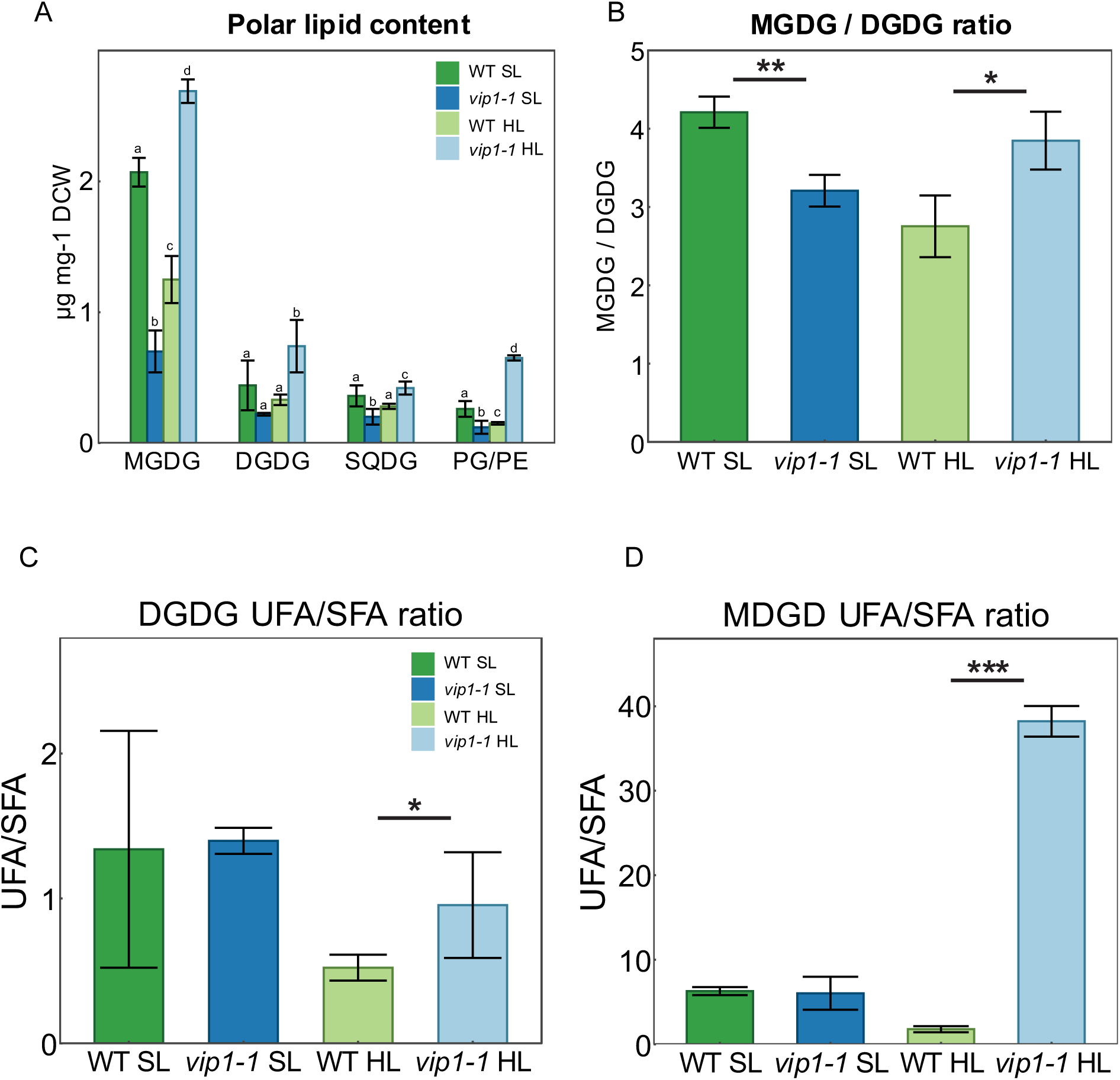
A) Quantification of polar lipid content, B) MGDG/DGDG ratio and C) fatty acid profile characterization in WT and vip1-1 strains under SL and HL conditions. Letters or asterisks (*, ** or *** for p-value < 0.05, 0.01 or 0.001, respectively) are used to denote significant differences. Data normality was verified using the Shapiro test, and significance was calculated using the t-test. Error bars indicate mean ± SD.

Interestingly, the MGDG/DGDG ratio was lower in *vip1-1* than in WT under SL, but higher under HL (Figure 3B). This revealed that not only the levels of the individual polar lipids responded to HL in an opposite manner in the absence of PP-InsPs, but also the MGDG/DGDG ratio. This ratio has been reported to be dynamically adjusted by the cell to modify membrane fluidity in response to light or heat stress (Chen et al. 2006; Demé et al. 2014). In this way, a higher MGDG/DGDG ratio increases the thylakoid membrane fluidity, while lower MGDG/DGDG ratio is adopted under heat or HL stress to reduce thylakoid membrane fluidity. Additionally, we identified and quantified the different fatty acids (FA) present in each polar lipid fraction and found important changes in the unsaturation rates between strains (Supplementary Table I). Indeed, it was especially remarkable the differences found in MGDG and DGDG fractions between the two strains under HL conditions (Figure 3C and D). While WT downregulates UFA /SFA, *vip1-1* significantly upregulates this ratio reaching 1.6 fold in the case of DGDG (Figure 3C) and 11.6 fold in the case of MGDG (Figure 3D). Major differences were observed in C16:3; C16:4 and C18:3 which are mainly considered fatty acids present in the thylakoidal membranes (Supplementary Table I).

The MGDG profile from cells grown under SL was composed of similar saturated FA, less mono-unsaturated FA and more poly-unsaturated FA in *vip1-1* compared to the WT strain (0.51% more, 16.19% less and 15.67% more, respectively), depicting a higher desaturase activity in the PP-InsPs depleted strain. Indeed, when cells were grown under HL, the difference between strains increased to 33.41% less saturated FA, 8.78% less mono-unsaturated FA and 42.20% more poly-unsaturated FA in *vip1-1* compared to WT. This increased unsaturation rate as a response to HL in the *vip1-1* strain was observed in all the polar lipids studied (Supplementary Table I).

### 3.4 Gene Onthology revealed an important enrichment in photosynthesis and lipid biosynthesis

Following this lipid analysis, we conducted RNA-seq analysis under the studied conditions to gain deeper insight into the role of PP-InsPs in the regulation of HL response. First, we performed a Gene Ontology (GO) enrichment analysis of the activated genes under SL and HL conditions in *vip1-1* compared to the WT (Figure 4A and B). Under SL, the three most significantly enriched GO terms were related to photosynthesis (Figure 4A), but we also observed an enrichment in photosynthesis, translation, protein folding and protein localization GO terms under HL (Figure 4B).

**Figure 4:**
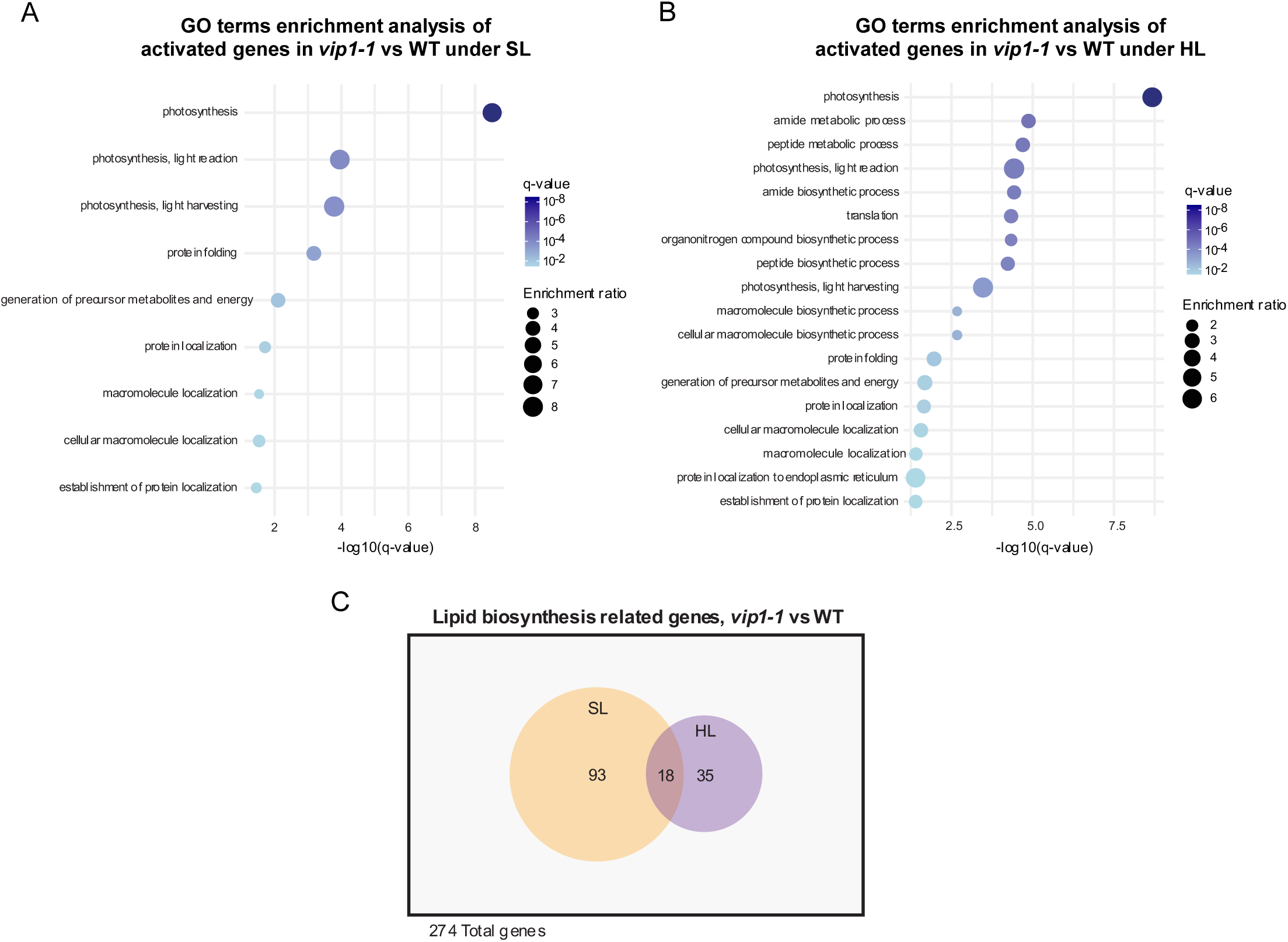
Bubble plots of enriched GO terms of activated genes in vip1-1 vs WT, A) under SL and B) under HL conditions. C) Venn diagram representing the number of differentially expressed genes related to lipid biosynthesis in vip1-1 vs WT under SL and HL conditions.

Building on the previously described lipid analysis results, we specifically analyzed the expression of genes associated with the lipid biosynthetic process (GO term GO:0008610). Of the 274 genes assigned to this GO category in *C. reinhardtii*, 93 were significantly differentially expressed genes in SL, 35 in HL and 18 in both conditions (Figure 4C). These findings led us to further investigate genes directly involved in FA metabolism. Firstly, we selected genes that were differentially expressed under either SL or HL (adjusted *p*-value < 0.05) and classified them into three categories: fatty acid synthases, glycerolipid synthases and fatty acid desaturases (Figure 5). In particular, the genes composing the glycerolipid synthases category have the following function: *sqd1* and *sqd2* are responsible for SQDG synthesis, *lat1/lpt1* is responsible for MGMG recycling into MGDG, *dgd1* is responsible for DGDG synthesis and *pgps3* is responsible for PG synthesis (Li-Beisson, Beisson and Riekhof, 2015; Iwai et al. 2021). Secondly, their expression patterns were visualized in a heatmap using row-wise Z-score normalization (Figure 5, Supplementary Table II). Under SL, 8 out of 9 genes involved in FA synthesis, 4 out of 5 genes involved in glycerolipid synthesis and 7 out of 9 genes involved in FA desaturation were upregulated in *vip1-1* compared to WT (Figure 5). Only 3 genes were significantly downregulated, *mct1*, *fad5-like2* and *fad6*; and *dgd1* was equally expressed in both strains (Figure 5, Supplementary Table II). On the other hand, the Malonyl-CoA:ACP transacylase *mct1* and the plastic desaturase *fad6* gene (Sato et al., 1997) were upregulated in the WT under SL conditions compared to *vip1-1* under the same conditions. These results show important differences in the expression of lipid biosynthesis related genes under SL and suggest the regulatory role of PP-InsPs in this metabolic process. In contrast, less differences were observed under HL conditions, as only *sqd1, sqd2, fad7* and *fad9* kept significantly upregulated in the *vip1-1* strain compared to the WT. Interestingly, the WT strain highly upregulated the expression of *dgd1* under these conditions, while the *vip1-1* strain did not (Figure 5). This is consistent with the upregulated levels of MGDG/DGDG ratio in *vip1-1* cells as *dgd1* catalyzes the conversion of MGDG into DGDG and controls their levels under stress conditions (Légeret et al., 2016). *Fad5 like 2* was also upregulated in the WT in contrast to *vip1-1* under HL (Figure 5). These results show a tight relationship between PP-InsPs and lipid metabolism at the gene expression, lipid accumulation and functional levels that need to be evaluated for future biotechnological applications of green organisms.

**Figure 5:**
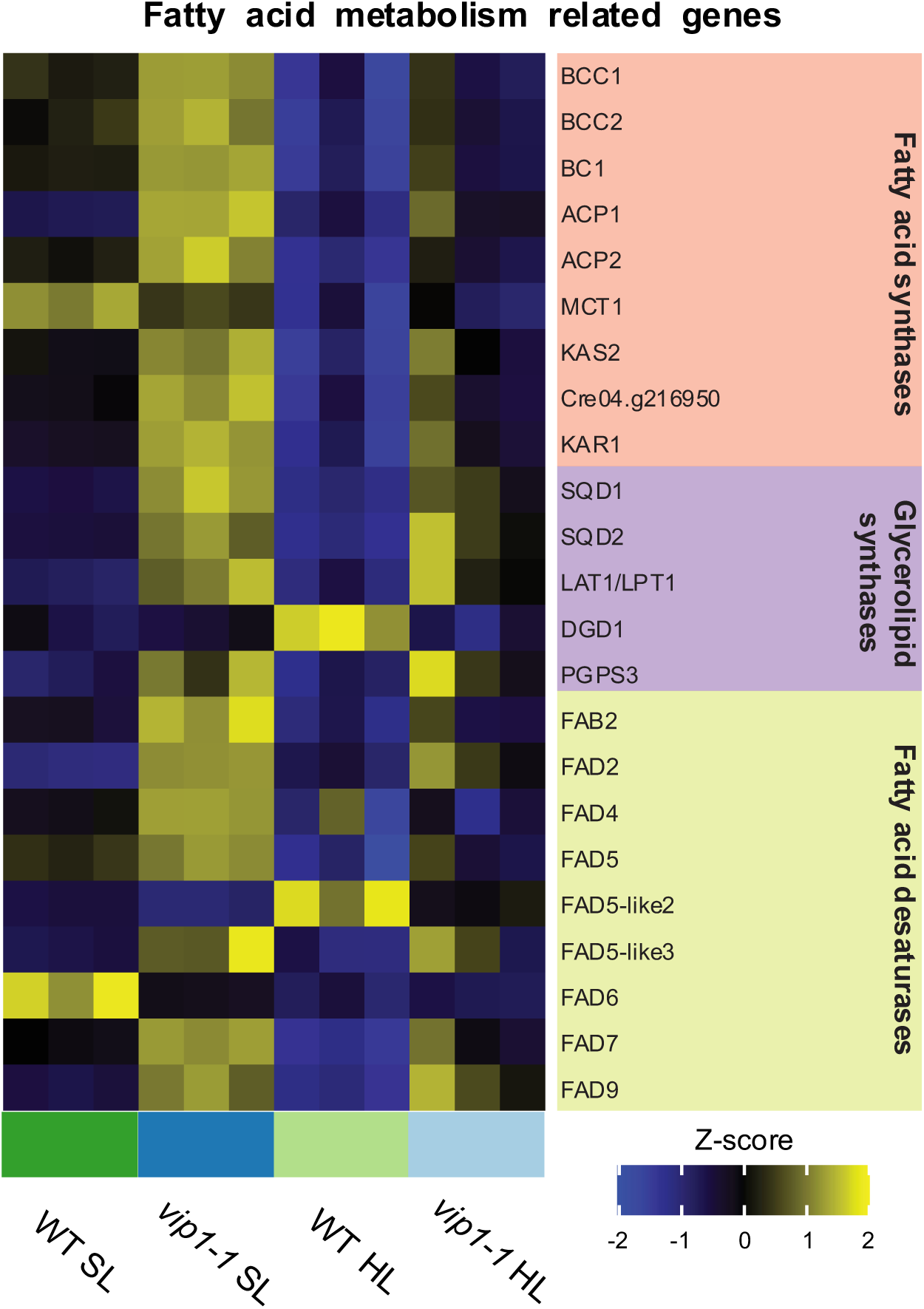
Heatmap depicting the differences in expression of genes involved in fatty acid synthesis, polar lipid synthesis and desaturations. Each gene is differentially expressed (adjusted p-value < 0.05, calculated using limma with BH adjusting method) in vip1-1 vs WT in at least one condition (SL or HL).

### 3.5 PP-InsPs are part of the regulatory mechanisms that control photosynthetic gene expression

Depletion of PP-InsPs resulted in alterations in the NADPH/ATP pool, a reduction in active PSII under high light (HL) conditions (Figure 1B), and changes in thylakoid lipid composition (Figure 3A, B and C), we hypothesized that these changes might be accompanied by modifications in the expression of genes involved in photosynthesis and light acclimation. After RNAseq data analysis by GO, we confirmed an enrichment of this process under SL and HL conditions (Figure 4A and B) that we further analyzed. We filtered photosynthesis-related differentially expressed genes (PhANGs) using a minimum fold-change threshold of 1.75 and an adjusted *p*-value of < 0.05. Under SL conditions, numerous genes encoding components of the photosystems (PS) and light-harvesting complexes (LHCs) were upregulated in the *vip1-1* strain compared to the WT (Figure 6A, Supplementary Table III). Among them, we found two members of the PsbP-like family, some *Conserved in the Green Lineage* (CGL) genes, and *pgr5*, a key regulator of cyclic electron flow (Alric, 2014). In contrast, the three light harvesting complexes stress-related (LHCSR) genes present in *C. reinhardtii* (*lhcsr1*, *lhcsr3.1* and *lhcsr3.2*), which mediate non-photochemical quenching under HL stress (Girolomoni et al. 2019; Wilson et al. 2023), were significantly downregulated in the mutant (Figure 6A, Supplementary Table III). Under HL conditions, the strong activation observed in SL was partially alleviated. Nevertheless, we found upregulations in the expression of four genes encoding LHC subunits, *PsaH*, *PSBP5* and *CGLD14* genes in *vip1-1* compared to WT (Figure 6A, Supplementary Table III). Remarkably, the expression of *lhcsr3.1* and *lhcsr3.2* genes in *vip1-1* reached the level in WT, and the *lhcsr1* gene was upregulated in the mutant under HL conditions. *pgr5* and *pgrl1* which are involved in cyclic electron flow regulation (Alric, 2014) kept upregulated in the *vip1-1* strain under HL (Figure 5A, Supplementary Table III).

**Figure 6:**
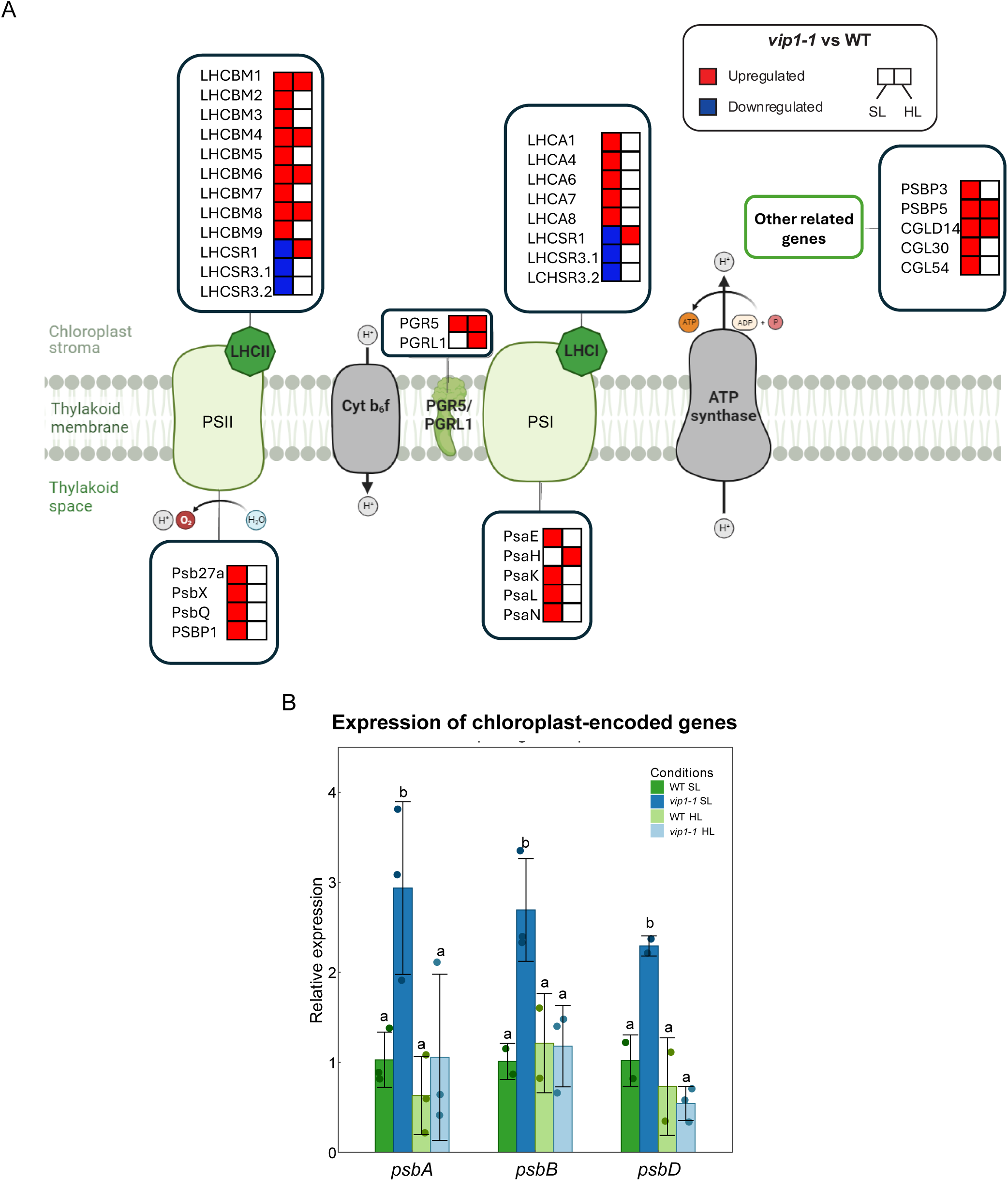
A) Expression of photosynthetic associated nuclear genes. Only genes with significant differences in at least one condition are represented (adjusted pvalue < 0.05, calculated using limma with BH adjusting method). B) Expression of chloroplast-encoded genes. Letters are used to denote significant differences. Data normality was verified using the Shapiro test, and significance was calculated using the t-test. Error bars indicate mean ± SD.

As there are several chloroplast-encoded genes related to photosynthesis, we also analyzed and compared their expression patterns with related nuclear genes. We quantified the relative expression levels of *psbA*, *psbB*, and *psbD*, using *cblp* as the reference housekeeping gene (von Kampen et al., 1994). The expression patterns of these genes were similar under both SL and HL conditions in WT. In contrast, *vip1-1* accumulated significantly higher transcript levels than the WT under SL, while under HL no significant differences were detected between strains (Figure 6B). However, the expression of chloroplast-encoded genes (Figure 6B) is similar to the response we found in the expression levels of PhANGs (Figure 6A). This can be indicative of a coordination on the photosynthetic gene expression, PhANGs and chloroplast encoded, regulated by PP-InsPs under control conditions.

### 3.6 PP-InsPs regulate CBB cycle at transcriptional and metabolic level

Following the activation of photosynthesis-associated genes in the *vip1-1* under both SL and HL conditions, we conducted a metabolomic analysis to determine whether increased transcript levels were accompanied by changes in the accumulation of Calvin-Benson-Bassham (CBB) cycle intermediates. To obtain a multiomic perspective, we integrated the metabolic profiles with the RNA-seq data of CBB-related enzymes (Figure 7). Under SL conditions, *vip1-1* showed an overaccumulation of 10 out of the 11 CBB-related metabolites detected compared to WT, with the exception of 3-phosphoglycerate (Figure 7, Supplementary Table IV). Additionally, transcriptional analysis revealed that *pgk1*, *tpic1*, *fbp2*, *rpi1* and *rpe2* were activated in *vip1-1* in the same conditions. These genes encode enzymes responsible for the phosphorylation of 3-phosphoglycerate (*pgk1)*, isomerization of glyceraldehyde 3-phosphate and dihydroxyacetone phosphate (*tpic1*), dephosphorylation of fructose 1,6-bisphosphate (*fbp2*), and the isomerization of ribose 5-phosphate, xylulose 5-phosphate and ribulose 5-phosphate (*rpi1* and *rpe2*). These results suggest a localized activation of the CBB cycle enzymes under SL in response to PP-InsPs depletion, leading to the accumulation of C in form of CBB intermediates. Under HL conditions, six metabolites remained overaccumulated in *vip1-1* compared to WT (ribulose 1,5-phosphate, dihydroxyacetate phosphate, glyceraldehyde 3-phosphate, fructose 1,6-bisphosphate, erythrose 4-phosphate and ribose 5-phosphate), while the levels of 4 of them were similar to WT levels: ribulose 5-phosphate, fructose 6-phosphate, sedoheptulose 7-phosphate and xylulose 5-phosphate (Figure 7, Supplementary Table IV). In addition, we observed important differences at the transcriptomic level. Indeed, only *rpi1* and *fbp2* were expressed in *vip1-1* at similar levels of WT, while *prk1* (phosphoribulokinase), *rpe2* (xylulose 5-phosphate and ribulose 5-phosphate isomerase), *fba3* (fructose bisphosphate aldolase), *rbcs1* and *rbcs2* (RuBisCO small subunits) were activated in the mutant strain under HL. Therefore, our results provide evidence of an increased transcription of photosynthetic machinery and CBB cycle enzymes, together with an accumulation of CBB intermediates in *vip1-1*, both in SL and HL (Figures 6A, 6B, 7, Supplementary Table III, IV). These revealed an important role for PP-InsPs in the regulation of carbon assimilation at transcriptional and metabolic levels.

**Figure 7:**
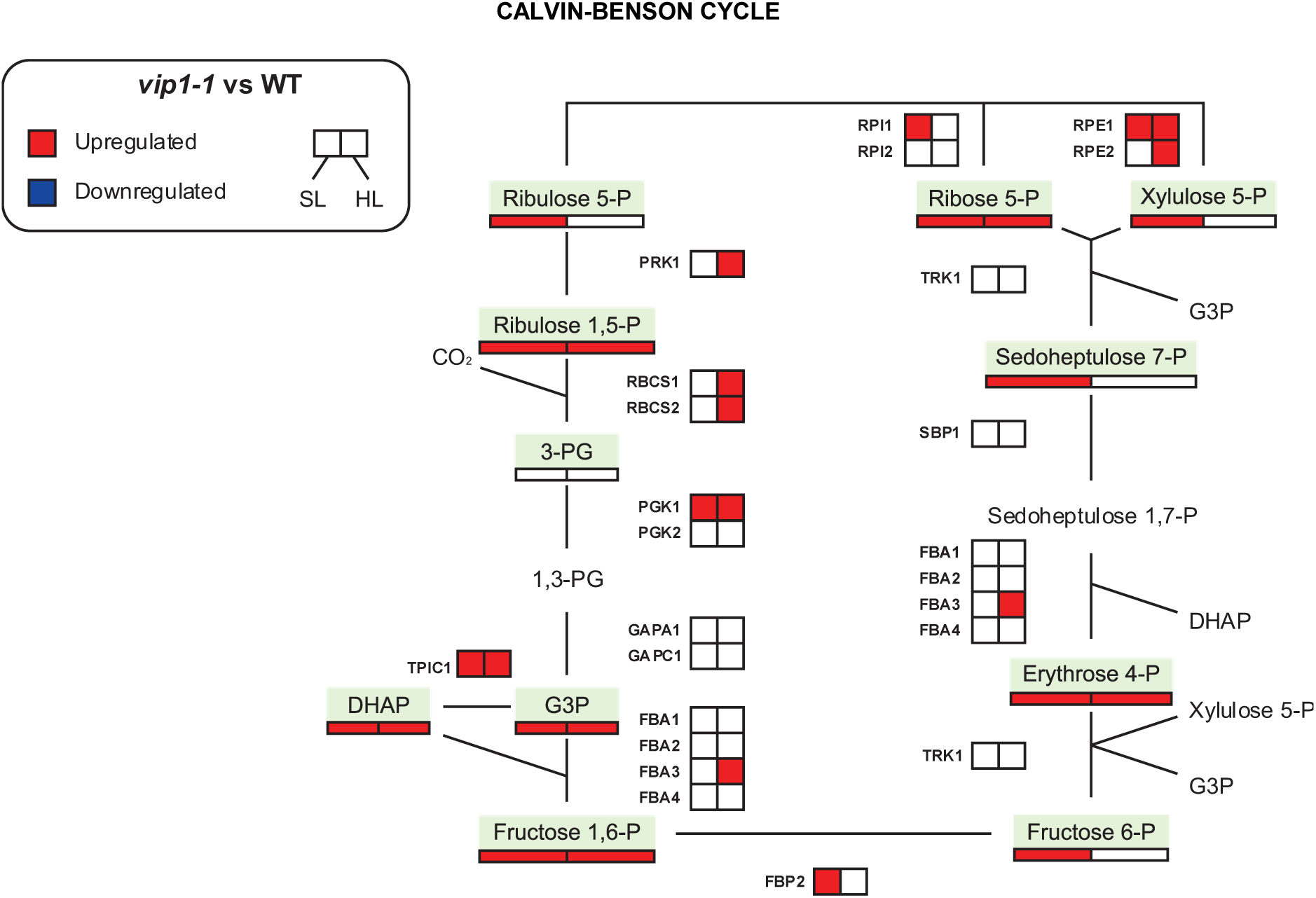
Gene expression and metabolite accumulation of Calvin-Benson intermediates and enzymes. Genes and metabolites are considered differentially expressed or accumulated when the differences in transcript or metabolite accumulation between vip1- 1 and WT strains greater than 1.75-fold and the adjusted p-value less than 0.10. Significance was calculated using the Wilcoxon rank-sum test for metabolite accumulation and limma with BH adjusting method for gene expression. Non-evaluated metabolites are displayed without a green box.

### 3.7 Chlorophyll synthesis regulation is impaired in *vip1-1* mutant

In photosynthetic organisms, PS core subunits and LHC subunits are required to bind chlorophylls to correctly organize into supercomplexes. Indeed, the process of assembly and disassembly is tightly regulated to avoid toxicity derived from free chlorophylls (Wang and Grimm, 2021; Willows, Lagarias and Duanmu, 2023). In this way, chlorophyll synthesis must be coordinated with the availability of chlorophyll binding elements (Wang and Grimm, 2015). Given that, we found a widespread gene expression activation of LHC and PS subunits in the PP-InsPs depleted strain, we next performed chlorophyll measurements and analyzed gene expression of the tetrapyrrole biosynthetic pathway to determine whether the chlorophyll synthesis was altered too. Notably, the chlorophyll a/b ratio (Figure 8 A) was lower in the mutant strain under HL. While chlorophyll a can be found in the LHCs, PSI and PSII, the chlorophyll b is only located at the LHCs. This way, the chlorophyll a/b ratio can be used to estimate the antenna size (Dupuis et al. 2025). Therefore, the significant decrease in the chlorophyll a/b ratio in the mutant strain under HL suggests *vip1-1* has an increased antenna size compared to WT.

**Figure 8:**
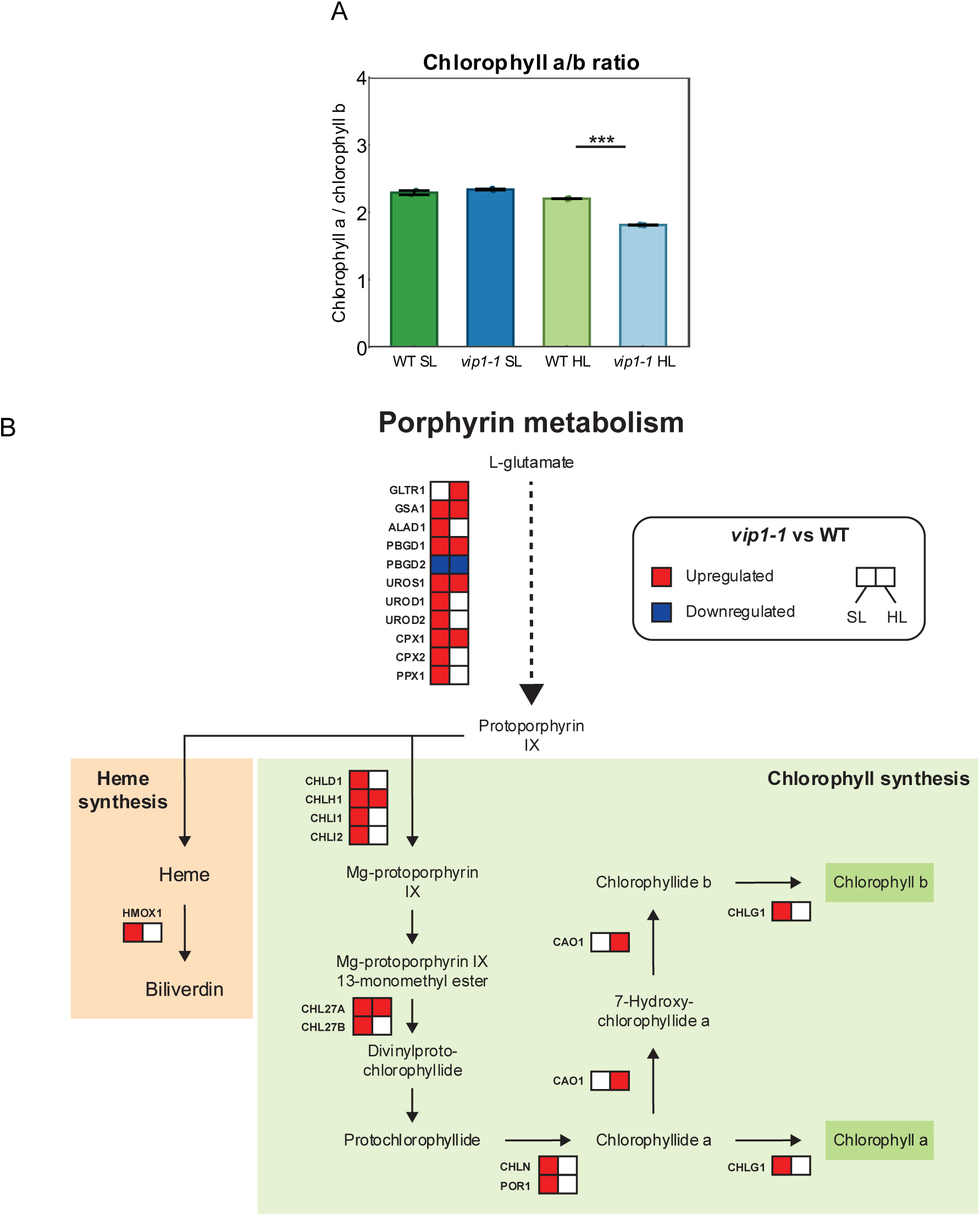
A) Chlorophyll a/b ratio, calculated as chlorophyll a / chlorophyll b for each replicate. Asterisks (*, ** or *** for p-value < 0.05, 0.01 or 0.001, respectively) are used to denote significant differences Data normality was verified using the Shapiro test, and significance was calculated using the t-test. Error bars indicate mean ± SD. B) Expression of nuclear genes involved in the porphyrin biosynthetic route and CHLN (chloroplast-encoded). Genes are considered diferentially expressed if adjusted p-value < 0.05, calculated using limma with BH adjusting method. CHLN significance was calculated using t-test.

We next examined the expression of the genes involved in the tetrapyrrole biosynthetic pathway. Through a series of enzymatic reactions, L-glutamate is converted into protoporphyrin IX, which represents the branching point where heme and chlorophyll biosynthetic routes diverge (Willows, Lagarias and Duanmu, 2023). In the shared part of the pathway, we observed significantly higher accumulation of transcripts in the mutant under SL conditions, which was partially maintained under HL conditions (Figure 8B). Additionally, the heme oxidase *hmox1* was found to be upregulated in SL conditions in the *vip1-1* strain. This enzyme is responsible for the synthesis of biliverdins, which are linear tetrapyrroles involved in photoprotection and the modulation of the magnesium chelatase subunit CHLH1 activity (Zhang et al. 2021). However, the most pronounced differences were observed in the chlorophyll-specific branch of the pathway. Under SL, *por1* and several other genes encoding magnesium chelatase subunits were found upregulated in *vip1-1* (*chld1*, *chlh1*, *chli1*, *chli2*, *chl27a*, *chl27b*, *chln* and *chlg1*) (Figure 8B). Interestingly, when cells were grown under HL, most magnesium chelatases differences were alleviated, but chlorophyllide a oxigenase (*cao1*) transcripts were found to be higher accumulated in the mutant strain. This gene is responsible for the conversion of chlorophyll a to chlorophyll b in *C. reinhardtii*, which is consistent with the lower chlorophyll a/b ratio found in the mutant under these HL conditions (Figure 8A and B) (Bujaldon et al. 2017). Our results suggest that the depletion of PP-InsPs significantly upregulates transcripts related to photosynthesis and tetrapyrrole biosynthesis under SL conditions.

Aberrant carotenoids levels have been associated with mutants affected in antenna size (Kirst et al., 2012). In this regard, we also analyzed carotenoid levels to evaluate possible alterations which could additionally affect the photosynthetic apparatus structure or photoprotective capacity. We found that primary carotenoids levels (α-carotene, β-carotene and lutein) were similar between strains under both conditions tested (Supplementary Figure 2A). Nevertheless, we identified significant differences in the xanthophyll cycle. This cycle is a photoprotective mechanism that comprises violaxanthin, antheraxanthin and zeaxanthin, where violaxanthin can be de-epoxidized to produce antheraxanthin and zeaxanthin upon light stress, the latter being essential for non-photochemical quenching (NPQ) activity (Niyogi, Björkman and Grossman, 1997; Baroli et al. 2003). Our results revealed a higher de-epoxidation state of the xanthophyll cycle in the *vip1-1* strain compared to the WT under both SL and HL conditions (Supplementary Figure 2B). In this way, PP-InsPs are connected to the regulation of photoprotective mechanisms in *Chlamydomonas* cells.

### 3.8 GO and amioacid analysis show an upregulation of protein synthesis in *vip1-1*

Gene Ontology (GO) analysis of differentially expressed genes revealed that the biological processes of “protein folding” and “protein localization” were significantly enriched in the activated genes in *vip1-1* regardless of the light conditions (Figure 4A and B). Additionally, the protein translation was also significantly enriched in the mutant strain under HL (Figure 4B). Consistently, KEGG pathway enrichment analysis of activated genes in *vip1-1* indicated significant enrichment of “Protein processing in endoplasmic reticulum” pathway under both SL and HL conditions, and “Protein export” pathway under HL (Supplementary Table V). This information led us to further characterize these responses by selecting genes related to these processes that were differentially expressed under either SL or HL (adjusted *p*-value < 0.05). Expression levels were visualized using a heatmap, with clustering rows based on unweighted pair group method with arithmetic mean (UPGMA) and Pearson distance (Supplemental Figure 3).

In addition, given the alterations observed in the chloroplast lipid composition and photosynthesis-associated gene expression (Figures 3A, 3B, 6A and 6B), we analyzed the subcellular localization of the proteins encoded by each gene to assess whether the differential regulation occurred in a compartment-dependent manner (Supplemental Figure 3). We identified 179 differentially expressed genes associated with protein synthesis. Of these, 50 of the proteins encoded by these genes were targeted to the chloroplast, 47 to the mitochondrion, 8 to the secretory pathway and 74 to the cytosol (Supplementary Table VI). Most of these genes encode ribosomal subunits or translation factors, and their expression patterns were found to be dependent on the subcellular location of the gene product. Genes encoding chloroplast-targeted proteins clustered closely, showing a moderate increase in their expression in the *vip1-1* mutant strain under both SL and HL conditions (Supplemental Figure 3). Furthermore, genes encoding mitochondrion-targeted proteins also clustered closely, showing a more pronounced upregulation in their gene expression in the mutant strain under both light conditions (Supplemental Figure 3). Given the increased transcription of genes encoding chloroplast and mitochondrion-localized proteins, we hypothesized that the expression of the importing complexes of these organelles should be higher in *vip1-1* too. In this context, we found an increased expression of endoplasmic reticulum (*erd2a*, *sec62*, *sec61b*, *srp72* and *srp1*), Golgi (*cog7* and *copb2*), chloroplast (*ftsy* and *alb3.1*) and mitochondrion (*tom7*, *tim17* and *tim50*) translocases, which are significantly upregulated both in SL and HL conditions in *vip1-1* (Supplementary Table VII). These results provide evidence of an induced protein synthesis in the context of PP-InsPs depletion in the *vip1-1* strain.

To corroborate these findings, we analyzed the accumulation of amino acids in the WT and mutant strain under SL and HL conditions (Figure 9). Under SL, glutamine and arginine levels were greatly reduced in the *vip1-1* strain (Supplementary table VIII). In contrast, alanine, valine, methionine and glutamate were significantly accumulated in the mutant under SL conditions. However, under HL conditions, arginine levels were further reduced by 2.49-fold in the *vip1-1* strain compared to the WT while the glutamine levels difference was alleviated. Remarkably, under this stress we found a significant decrease in the levels of glutamate and in the cluster formed by leucine, isoleucine, phenylalanine, aspartate and tryptophan in the *vip1-1* strain compared to the WT (Figure 9, Supplementary Table VIII). This decrease in amino acid levels is consistent with the higher expression levels of protein synthesis-related genes observed in *vip1-1* specially under HL (Supplemental Figure 3), which suggests a modulatory role of the PP-InsPs in the integration of protein synthesis and environmental cues. Therefore, we observed not only an induction of structural carbon sinks in the form of chloroplast lipids, but also an increased allocation of carbon toward protein synthesis.

**Figure 9:**
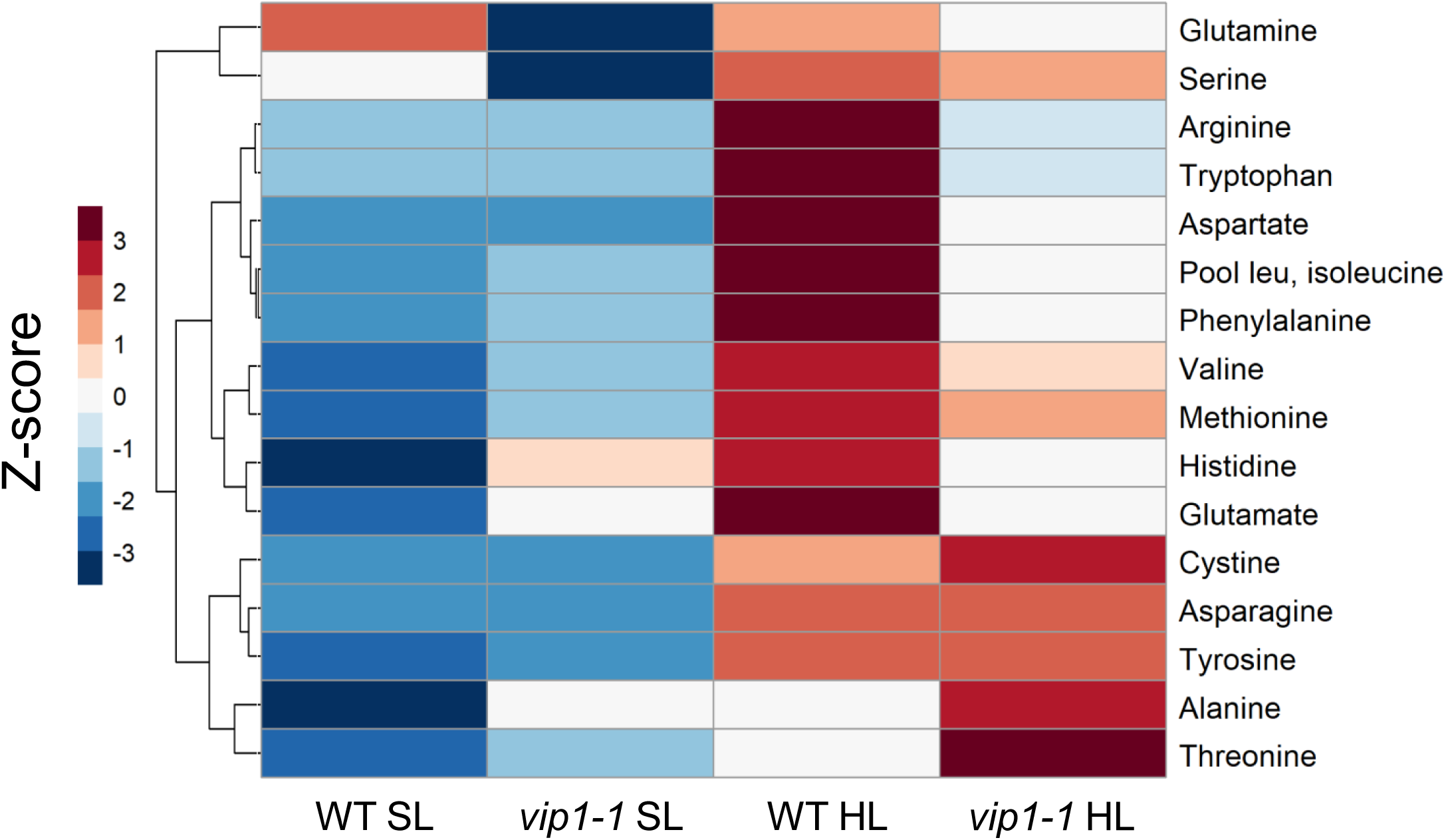
Heatmap of amino acid accumulation in WT and vip1-1 under SL and HL conditions. Rows were clustered using complete-linkage method with euclidean distance.

## 4 Discussion

Photosynthetic organisms need to balance light absorption and carbon fixation in accordance with the light intensity they are exposed. Fine-tuning of photoprotective mechanisms helps them thrive even when light irradiance is excessive. Here, we show that in *C. reinhardtii*, inositol pyrophosphates (PP-InsPs) are multifaceted molecules that take part in adjusting photosynthetic activity (Figures 1 and 6), carbon fixation (Figure 7) and structural carbon pools (Figures 2, 3, 5, 8 and 9) to optimize energy production and carbon storage in accordance with light intensity.

Inositol polyphosphates have been associated to photosynthesis regulation, through both protein phosphorylation (Couso et al. 2021) and integration of nutrient availability like nitrogen or CO_2_ (Morales-Pineda et al. 2022; Bedera-García et al. 2025). In this work, we found that PP-InsPs deficient mutant *vip1-1* is sensitive to high light (Figure 1A and B). However, little is known about the relationship between these highly phosphorylated molecules and light stress. The results on TL analysis suggest that there is a significant alteration in photosynthesis associated to *vip1-1* mutation (Figure 1B). Under SL, we observed differences in the AG band compared to WT indicating less availability of NADPH and/or ATP (Mellvig and Tillberg 1986; Palmqvist et al. 1986; Sundblad et al. 1986; Krieger et al. 1998). Currently, there is no reported evidences in the direct connection between PP-InsPs and NADPH but impairment in PP-InsPs synthesis have previously been connected to ATP levels in mice or yeasts (Szijgyarto et al., 2011). In addition, InsP_7_ stimulates polyP synthesis based on ATPase activity (Wild et al., 2016) that influences ATP availability. Although we cannot directly link the deregulation of photosynthesis in our results to ATP availability in *vip1-1* cells, our data suggest that PP-InsPs play an important role in the regulation of energy levels in green cells. Our current TL results also indicate that there are less active PSII complexes in the mutant under HL conditions than in WT. This can be connected to the aberrant phosphorylation patterns of PSII proteins in *vip1-1* (Couso et al., 2021). Taking together, these results may indicate that PP-InsPs are likely involved in the correct regulation and photoprotection of PSII activity upon light stress. We also observed that InsP_7_ and InsP_8_ were upregulated under HL conditions in the WT (Figure 1C and D), indicating a stress-induced response in inositol pyrophosphate biosynthesis that has not been described before in green organisms.

Lipids and starch are major carbon and storage compounds in green microalgae that compete for the carbon use (Li et al., 2010). In Couso et al., 2016, we found *vip1-1* overaccumulates lipids in the form of TAGs under N deficiency and rapamycin treatment. In the case of high light conditions, we have found an important accumulation of starch granules in the WT that was not seen in *vip1-1* (Figure 2A). Biochemical determination confirmed a significant increase of starch in WT cells under HL that was not observed in *vip1-1* cells content that remained the same levels as in SL conditions (Figure 2B). Besides this, we analyzed total FA (Figure 2C) and TAGs (Figure 2D) and found increased levels in *vip1-1* compared to WT under both SL and HL conditions. These data add to our previous results on the important role of PP-InsPs in carbon partitioning as regulatory elements in central carbon flow.

Furthermore, we also evaluated the implications of PP-InsPs in the polar lipids, especially lipids from the thylakoid membranes to see if there was a connection between lipid synthesis and deregulation of photosynthesis in *vip1-1*. The thylakoid membranes are mainly formed by monogalactosyldiacylglycerol (MGDG), digalactosyldiacylglycerol (DGDG), sulfoquinovosyldiacylglycerol (SQDG) and phosphatidylglycerol (PG). These glycerolipids have a basic structural function in the chloroplast, but also present a stabilizing function in several protein complexes typical of photosynthetic organisms such as PSI, PSII, or Cyt *b6f* (Kobayashi, Endo and Wada, 2016). We have found *vip1-1* had significantly lower accumulation of MGDG, SQDG and PG/PE pool under SL, but significantly overacumulated all 4 major glycerolipids under HL stress (Figure 3A) compared to WT. Total fatty acid (FA) profile revealed a widespread accumulation of different FA species as a result of PP-InsPs depletion both at SL and HL conditions (Supplemental Figure 2). Together, the data suggest that PP-InsPs depletion drives cellular fatty acid accumulation and influences lipid regulation specifically within chloroplasts in a light-dependent manner, highlighting a possible role for PP-InsPs in chloroplast lipid management. MGDG/DGDG ratio was also disrupted as a consequence of PP-InsPs depletion (Figure 3B). This ratio is adjusted by photosynthetic organisms to control membrane fluidity to achieve HL and heat tolerance, as DGDG increases the membrane stacking due to the extra galactose present in its structure (Chen et al. 2006, Demé et al. 2014). We found that WT diminishes thylakoid membrane fluidity in response to HL by decreasing MGDG/DGDG ratio, while *vip1-1* cannot modify this ratio to the same extent. Therefore, we propose that PP-InsPs are involved in the regulation of not only the carbon partitioning of storage compounds, as reported previously in *C. reinhardtii* (Couso et al. 2016), but also the homeostasis of thylakoid lipids biosynthesis, which directly affects the photosynthetic process by controlling membrane composition, structure and fluidity.

Our transcriptomic analysis revealed increased expression of FA desaturases in the absence of PP-InsPs, both under SL and HL conditions, which is consistent with alterations observed in the total FA profile (Supplemental Figure 1 and Supplementary Table II). In fact, *vip1-1* consistently exhibited higher FA unsaturation rates than WT under HL conditions, with remarked differences found specially in MGDG unsaturation rates (MGDG unsaturated FA / saturated FA ratio of 1.78 in WT cells and 38.21 in *vip1-1* cells) (Supplementary Table I). Plastidial FA desaturation requires both the FA desaturases and reducing power provided by ferredoxin through ferredoxin-NADP oxidoreductase (Shanklin and Cahoon, 1998). FA desaturase transcripts were elevated in *vip1-1* under both at SL and HL, but substantial increases in unsaturation rates of plastid lipids were only observed in HL (Figure 4, Supplementary table II). This suggests that availability of reducing power in *vip1-1* under SL might be insufficient to perform these unsaturations, whereas under HL conditions the excess excitation likely provides enough reducing power to drive these reactions. Thermoluminescence data obtained under SL conditions (Figure 1B upper pannel) indicated lack of NADPH and/or ATP in the mutant. Reduced NADPH levels may affect fatty acid (FA) unsaturation, thereby linking PP-InsPs to the pool of energy-related molecules that regulate thylakoid membrane structure.

In addition, under SL we observed higher transcript levels of glycerolipid synthases in *vip1-1*, but lower levels of MGDG, SQDG and PG/PE pool as discussed above (Figure 3 and 5). We propose that these chloroplast-specific discrepancies between the expression of glycerolipid synthases and desaturases, and glycerolipid composition under SL reflect unfavorable stromal conditions for these enzymes. Together, these findings suggest that PP-InsPs may normally repress the expression of glycerolipid synthases and FA desaturases. Therefore, modifications in the abundance of these molecules could contribute to HL and stress acclimation through gene derepression and remodeling of thylakoid membranes. Reducing the levels of PP-InsPs (Figure 2C and 2D) could be part of the HL stress response in *C. reinhardtii*, which would explain why we observe a partial alleviation of the *vip1-1* phenotype in HL, as described in the results section (Figures 5, 6, 8B). In this context, removal of PP-InsPs could be used to prime the cell for HL and heat tolerance mechanisms by adjusting thylakoid membrane fluidity and structure. Nevertheless, additional regulatory layers such as PTMs, redox regulation or pH-dependent activity are likely required to fully activate these enzymes in the chloroplast, as reported for other chloroplastic processes and reactions (Lemaire, 2004; Ballotari et al. 2016; Grabsztunowicz et al. 2017; Camargo et al. 2021), including lipid synthases (Hernández and Cejudo, 2021 Morales-Sánchez et al. 2017). These studies support the idea that discrepancy between (i) transcriptional activation of desaturases and synthases and (ii) glycerolipid accumulation and composition might be a consequence of suboptimal plastid redox or pH conditions.

Thermoluminescence data indicate a significant lower AG band under SL and lower TL intensity under HL in *vip1-1* compared to WT which indicates less active PSII complexes (Figure 1B bottom pannel). In these conditions, we detected less differences on the expression of photosynthesis-associated nuclear genes (PhANGs) under HL compared to SL in *vip1-1* (Figure 6A) but some PSII components remained upregulated in the mutant. Interestingly, RNA levels of *lhcsr1* changed from downregulated under SL to upregulated levels under HL in *vip1-1* indicating that photoprotective mechanisms are highly upregulated in the mutant under light stress. Therefore, the combination of quenching mechanisms such as carotenoid de-epoxidation state (Supplemental Figure 2B) and *lhcsr1* increased transcription levels (Figure 6A) together with an alleviation of the differences in PSII subunits transcription (Figure 6A) can lead to the reduction of PSII active complexes as indicated by TL data in HL conditions (Figure 1B bottom panel).

The widespread transcriptional activation of the photosynthetic apparatus under SL in the mutant indicates a deregulatory response resulting from PP-InsPs depletion rather than a compensatory mechanism, as increased transcript abundance did not correlate with enhanced photosynthetic performance but instead pointed to abnormal NADPH/ATP pool as seen by a decrease in AG band (Figure 1B upper panel). We have previously discussed that PP-InsPs appear to act as fine-tuning molecules in stress acclimation by repressing the expression of genes encoding glycerolipid synthases and desaturases. Our transcriptomic analysis indicates that this PP-InsPs-dependent repression extends to both PhANGs and chloroplast-encoded genes too (Figures 6A and B). This coordinated regulation is consistent with the requirement for tight coupling between photosynthetic protein abundance and thylakoid membrane composition and fluidity to achieve optimal configurations in response to environmental cues such as light intensity or temperature. Thus, we propose that PP-InsPs form part of a repressive regulatory system that fine-tunes the expression of genes related to the photosynthetic process, integrating both from functional and structural aspects.

Additionally, under HL conditions, PP-InsPs depletion resulted in higher chlorophyll accumulation and larger estimated antenna size (Figure 8A). We suggest that this increase may be driven by the elevated levels of structural thylakoid lipids observed in the mutant strain under HL in order to maintain balanced stoichiometry between chlorophylls and membrane lipid components, required for stable photosynthetic complex assembly and function (Figure 3A) (Wang and Grimm, 2015; Kobayashi, Endo and Wada, 2016, Zhu et al. 2024). Supporting this idea, we found reduced levels of L-glutamate when chlorophyll synthesis is highly activated in *vip1-1*, which suggests that this amino acid is being used in the tetrapyrrole biosynthetic pathway (Figure 8B, 9; Supplementary Table VII).

Previous studies have identified small, phosphorylated molecules as key components of the chloroplast-nucleus communication system, including 3′-phosphoadenosine 5′-phosphate (PAP) or guanosine tetraphosphate (ppGpp) (Estavillo et al. 2011; Sugliani et al. 2016; Avilan et al. 2021; Romand et al. 2022). The functions of ppGpp have been extensively studied in different photosynthetic organisms like *Arabidopsis thaliana* (Sugliani et al. 2016; Romand et al. 2022), *Phaeodactylum tricornutum* (Avilan et al. 2021) and rice (Li et al. 2022). Although ppGpp has been detected in *C. reinhardtii* (Kasai et al. 2002; Bartoli et al. 2020), there is little information about its functional role in this green alga (Field, 2018). Based on our results, we believe that regulatory functions attributed to ppGpp in other photosynthetic organisms may be complemented by PP-InsPs in *C. reinhardtii.* Notably, ppGpp is known to participate in the regulation of choroplast gene expression in *Arabidopsis thaliana*, and in N starvation response in *Arabidopsis* and rice (Romand et al. 2022; Li et al. 2022). In line with this, we have found an extensive transcriptional reprogramming of both PhANGs and chloroplast-encoded genes caused by PP-InsPs depletion. Furthermore, we recently reported a compromised N starvation response as a consequence of PP-InsPs removal in *C. reinhardtii*, (Bedera-García et al. 2025). Together, these observations support the hypothesis that, in *C. reinhardtii* and possibly other photosynthetic organisms, PP-InsPs may fulfill regulatory roles attributed to ppGpp in other species. In addition to small phosphorylated nucleotides, tetrapyrrole intermediates such as heme, biliverdin and Mg-protoporphyrin IX have been identified as important mediators of chloroplast-nucleus communication where they can induce the expression of PhANGs (Strand et al. 2003; Zhang et al. 2021; Jan et al. 2022). In the PP-InsPs deficient strain *vip1-1*, we observed a misregulation of the tetrapyrrole biosynthetic pathway and increased PhANGs transcripts accumulation, as discussed above (Figures 6, 8B). These data suggest that nucleus-organelle communication might be compromised in the absence of PP-InsPs.

In line with the chloroplast-localized effects of PP-InsPs depletion discussed above, we found a mild transcriptional activation of the Calvin-Benson-Bassham (CBB) cycle under SL (Figure 7). In contrast to several other biological processes analyzed in this work, HL stress further increased the transcriptional activation of the CBB cycle genes in the PP-InsPs-depleted strain compared to the WT (Figure 7). Notably, metabolite analysis indicated the accumulation of most CBB cycle intermediates was higher in the absence of PP-InsPs under both SL and HL conditions, although we did observe partial alleviation of this accumulation under HL (Figure 7). This suggests that PP-InsPs repressive regulation extends to the process of carbon fixation and the accumulation of chloroplast-localized metabolic intermediates.

Our transcriptomic analysis also indicated that PP-InsPs depletion led to the activation of translocases and multiple biological processes related to protein synthesis and mobilization, such as protein folding, protein localization, protein processing in endoplasmic reticulum and protein export, regardless of light intensity (Supplementary Figure 3, Supplementary Table VII). PP-InsPs depletion is also associated with a significant reduction in free amino acid levels under HL conditions (Figure 9), and the previoulsy discussed accumulation of glycerolipids (Figure 3A). Taken together with previous reports showing that PP-InsPs depletion promotes triacylglycerol accumulation (Couso et al. 2016) and impairs starch accumulation (Bedera-García et al. 2025) under stress conditions, these findings support a central role for PP-InsPs in the regulation of carbon partitioning. Specifically, PP-InsPs appear to balance the allocation of carbon between storage pools (such as starch and triacylglycerols) and structural pools (such as proteins and thylakoidal glycerolipids) with HL response and possibly other stress responses. Therefore, we propose that PP-InsPs function as integrative signaling molecules that coordinate carbon pool utilization in response to both external and internal cues. This work reveals novel regulatory connections between inositol pyrophosphates and chloroplast-specific acclimation processes, providing useful information that may be used in future strategies to engineer resilient algal strains with minimal growth penalties in photobioreactors and other biotechnological applications.

## Acknowledgements

This research was supported by the Ministerio de Ciencia e Innovación TED2021-129409A-I00 and PID2022-136633OA-I00 grants awarded to IC. RBG was also awarded as a FPU22/00688 fellow by Ministerio de Educación.

## Author contributions

RBG and IC contributed to planning and experimental design; RBG, MEGG, LGHM, JMO, BP and IC performed experiments and data analysis; and RBG, JMO and IC wrote the manuscript.

## Data availability statement

The raw and processed RNA-seq data that support the findings of this study are openly available in the Gene Expression Omnibus (https://www.ncbi.nlm.nih.gov/geo/) repository and can be accessed with the Identifier GSE315948. The raw RNA-seq data are openly available at the Sequence Read Archive (https://www.ncbi.nlm.nih.gov/sra) and can be accessed with the identifier PRJNA1400007. The raw and processed metabolomic data are available from the authors upon request.

---

**Supplementary Figure 1:**
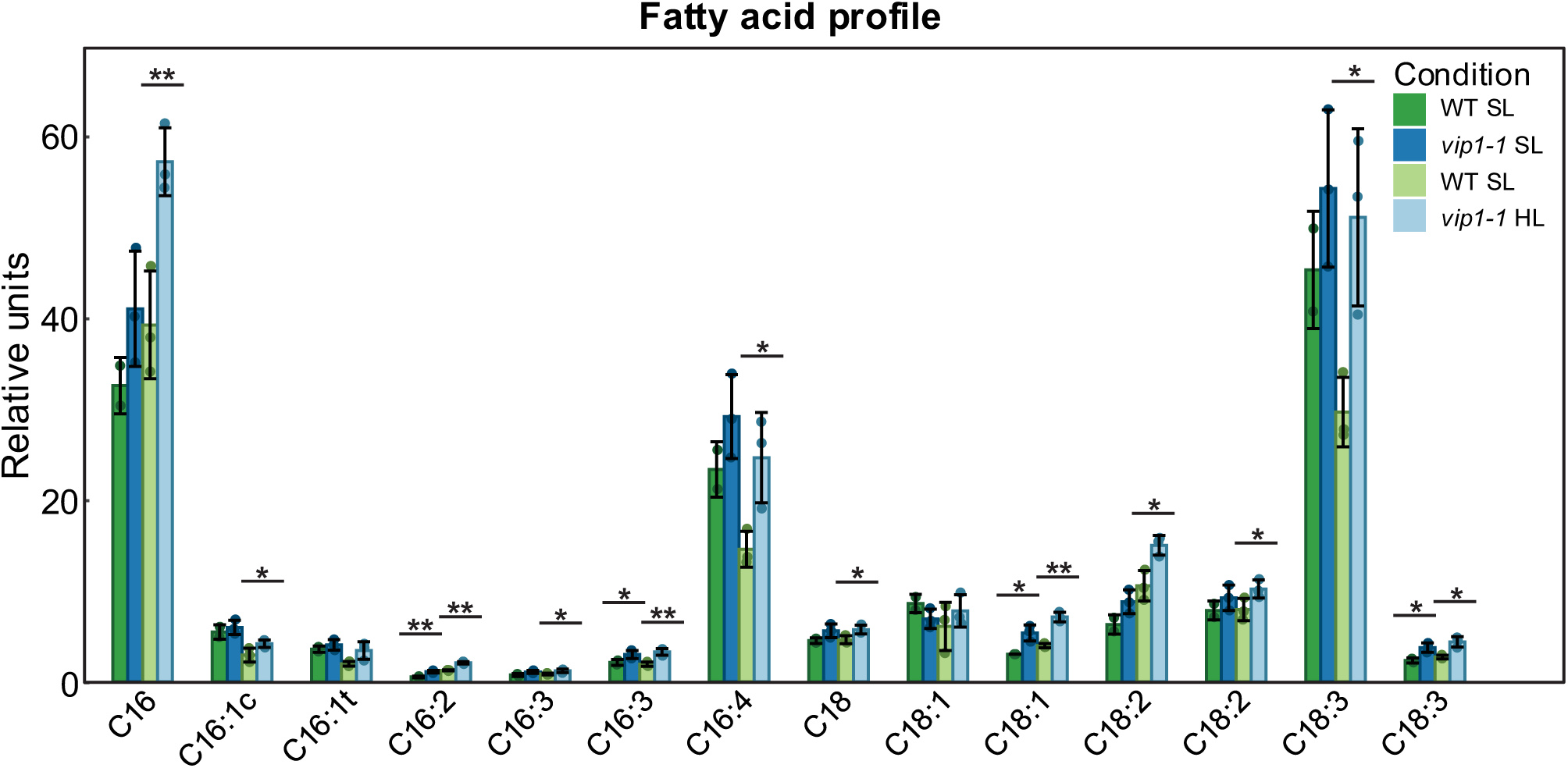
Total FA profile of WT and vip1-1 cells under SL and HL conditions. Asterisks (*, ** or *** for p-value < 0.05, 0.01 or 0.001, respectively) are used to denote significant differences. Data normality was verified using the Shapiro test, and significance was calculated using the t-test. Error bars indicate mean ± SD.

**Supplementary Figure 2:**
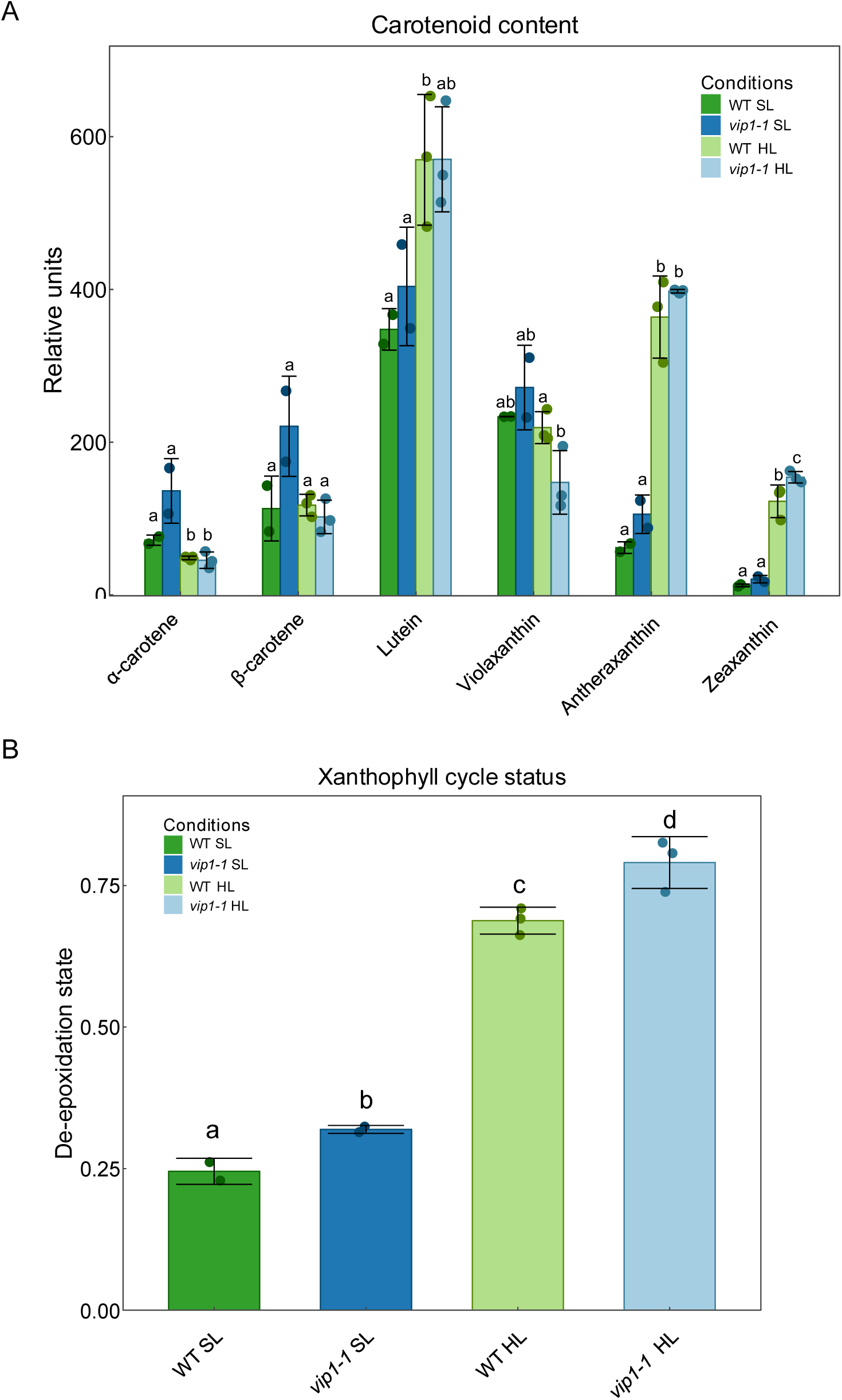
A) Carotenoid content of WT and vip1-1 strains under standard SL and HL conditions. B) De-epoxidation state of the xanthophyll cycle, calculated as: (Z + A)/(V + A + Z). Normality was verified using the Shapiro test, and significance was calculated using t-test. Letters indicate significant differences (p-value < 0.05)

**Supplementary Figure 3:**
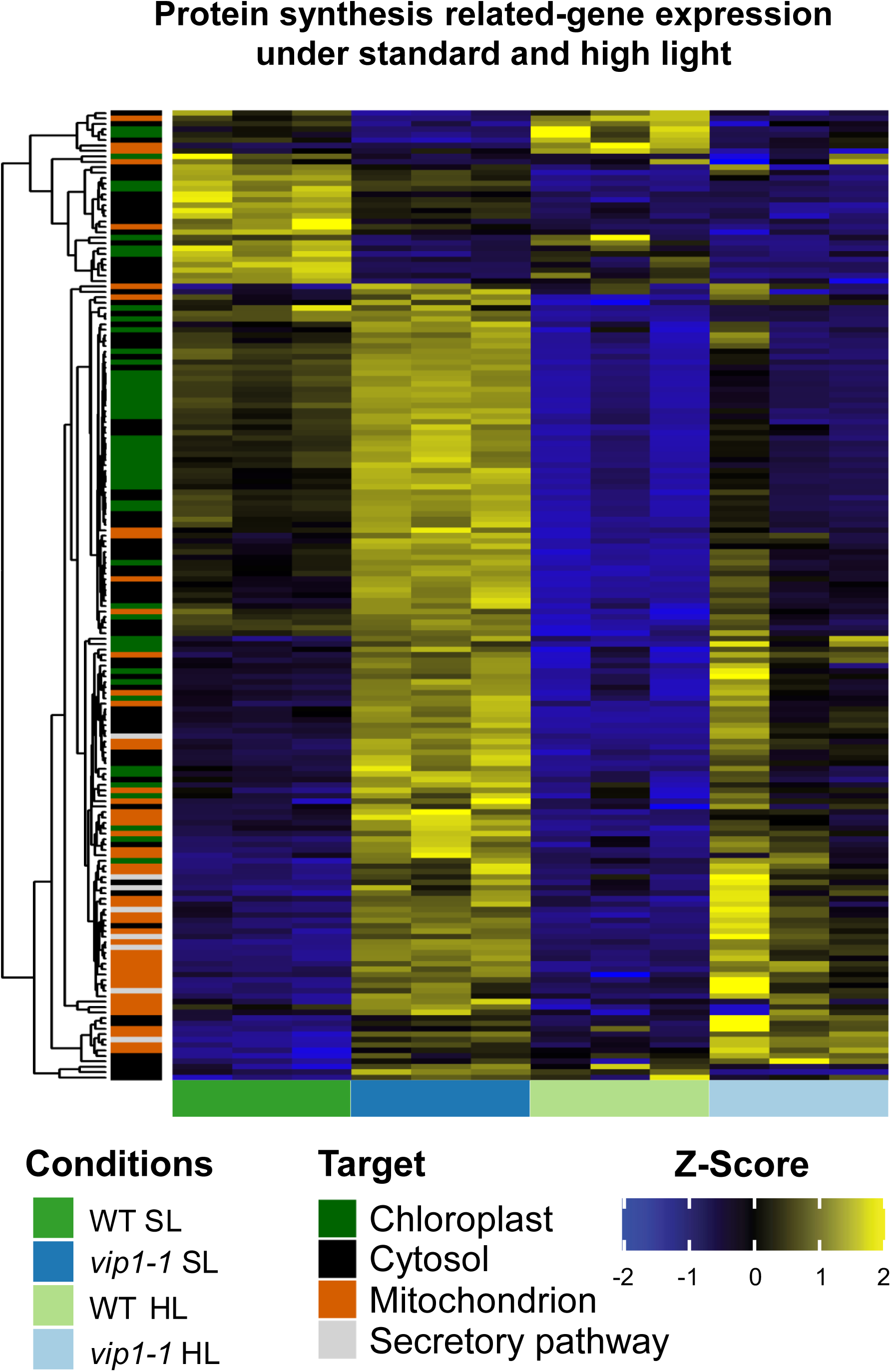
Heatmap of the gene expression of genes related to the protein synthesis process, and compartiment target of the protein encoded by each gene. Each gene is differentially expressed (adjusted p-value < 0.05, calculated using limma with BH adjusting method) in vip1-1 vs WT in at least one condition (SL or HL). BH djusting method) in *vip1-1* vs WT in at least one condition (SL or HL).

